# Independent innexin radiation shaped signaling in ctenophores

**DOI:** 10.1101/2022.10.11.511657

**Authors:** Jennifer Ortiz, Yuriy V. Bobkov, Melissa B. DeBiasse, Dorothy G Mitchell, Allison Edgar, Mark Q. Martindale, Anthony G. Moss, Leslie S. Babonis, Joseph F. Ryan

**Author notes:** These authors contributed equally (Jennifer Ortiz, Yuriy V. Bobkov).

## Abstract

Innexins facilitate cell-cell communication by forming gap junctions or non-junctional hemichannels, which play important roles in metabolic, chemical, ionic, and electrical coupling. The lack of knowledge regarding the evolution and role of these channels in ctenophores (comb jellies), the likely sister group to the rest of animals, represents a substantial gap in our understanding of the evolution of intercellular communication in animals. Here we identify and phylogenetically characterize the complete set of innexins of four ctenophores: *Mnemiopsis leidyi, Hormiphora californensis, Pleurobrachia bachei*, and *Beroe ovata*. Our phylogenetic analyses suggest that ctenophore innexins diversified independently from those of other animals and were established early in the emergence of ctenophores. We identified a four-innexin genomic cluster, which was present in the last common ancestor of these four species and has been largely maintained in these lineages. Evidence from correlated spatial and temporal gene expression of the *M. leidyi* innexin cluster suggest that this cluster has been maintained due to constraints related to gene regulation. We describe basic electrophysiological properties of putative ctenophore hemichannels from muscle cells using intracellular recording techniques, showing substantial overlap with the properties of bilaterian innexin channels. Together, our results suggest that the last common ancestor of animals had gap junctional channels also capable of forming functional innexin hemichannels, and that innexin genes have independently evolved in major lineages throughout Metazoa.

## INTRODUCTION

Ctenophores (comb jellies; Figure 1A) are marine animals defined by eight rows of ciliary paddles called comb rows. Ctenophores are capable of fast motor behaviors (e.g., ciliary reversal and tentacle retraction), have a number of neurosensory structures (e.g., nerve net, aboral organ, tentacles, sensory papillae), and possess a large suite of genes related to neural cell types and sensory information processing (Tamm 2014; Jager et al. 2011; Ryan et al. 2013). Nevertheless, ctenophore biology remains poorly understood, especially in relation to intercellular communication (Dunn et al. 2015). Genomic surveys in ctenophores looking for neurotransmitters and neurotransmitter pathways found in other animals have largely come up empty, and these results have formed the basis of the hypothesis that the ctenophore nervous system is a product of convergent evolution (Moroz et al. 2014). Phylogenomic evidence suggests that ctenophores are the sister to the rest of animals (Dunn et al. 2008; Figure 1B; Supplementary Table 1), and as such, are a key lineage for understanding the evolutionary history of animal nervous systems.

**Figure 1.**
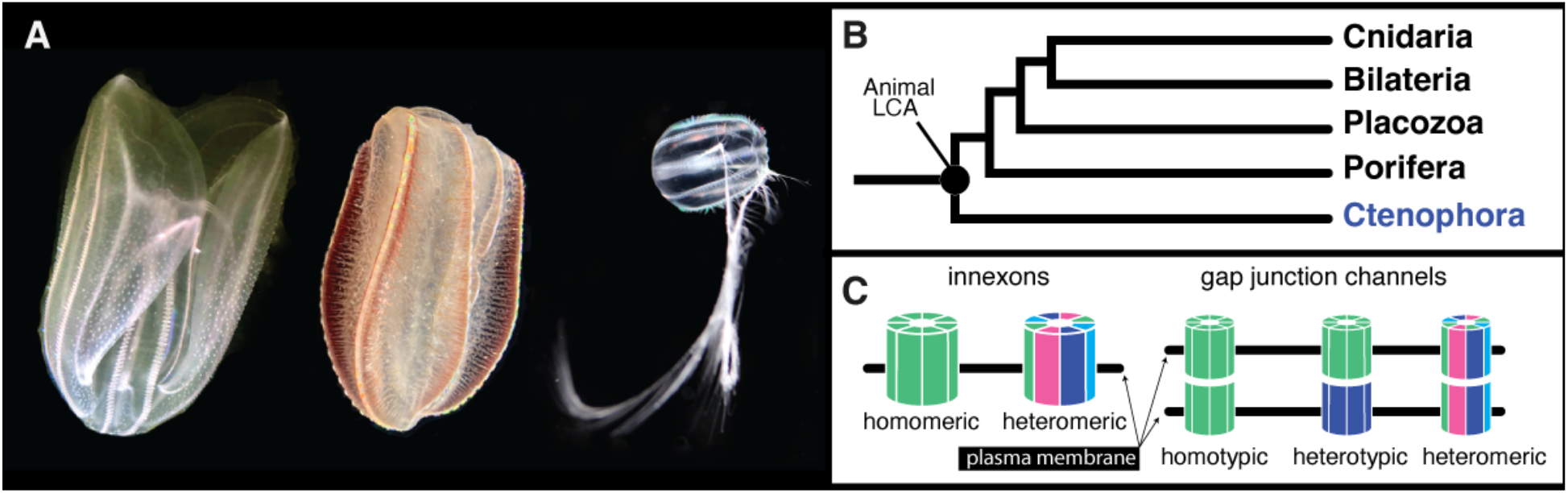
Ctenophores and innexins. (A) Three of the four ctenophore species in this study. From left to right: *Mnemiopsis leidyi, Beroe ovata*, and *Pleurobrachia bachei* (*Hormiphora californensis*, not pictured, is a tentaculate ctenophore with similar morphology to *Pleurobrachia*). (B) Based on phylogenomic evidence (Supplementary Table 1), Ctenophora is the sister group to the rest of animals. (C) Diagrammatic representation of potential subunit makeup of hemichannels and gap junctions (after Phelan and Starich 2001). Innexin subunits oligomerize to form a hemichannel. Hemichannels in adjacent cells can dock to form gap junctions. Hemichannels are either homomeric (composed of a single type of innexin) or heteromeric. Gap junctions are homotypic if hemichannels are identical, heterotypic if hemichannels are distinct, and heteromeric if hemichannels differ in subunit composition. *Mnemiopsis leidyi* photo by Arianna Rodriguez. Other photos by Joseph Ryan.

Gap junctions are assemblages of intercellular hemichannels that allow for the direct transfer of molecules and ions (up to ~1–3 kDa) between the cytoplasms of adjacent cells (Loewenstein 1966; Kanaporis et al. 2011; Oshima et al. 2013). In animals where gap junctions have been studied extensively, these channels have been detected joining virtually all cells in solid tissues (Goodenough and Paul, 2009). In the nervous system, gap junctions provide regulated pathways for the transfer of electrical signals (including bidirectional signaling), which often promote synchrony but can also provide a range of other functionality including inhibition and shunting of excitatory potentials (Vaughn and Haas, 2022). Furthermore, growing evidence indicate that electrical synapses could also be subject to activity dependent long-term plasticity (O’Brien and Bloomfield, 2018; Welzel and Schuster, 2018; Alcamí and Pereda, 2019).

In vertebrates, gap junctions are formed by hemichannels consisting of six subunits of connexin proteins (Yeager and Harris, 2007). Vertebrates also have channels made of pannexin proteins (usually eight subunits), which unlike connexins, do not form gap junctions but instead are able to connect the cytoplasm of cells directly to the extracellular environment (Dahl and Locovei, 2006; Sosinsky et al. 2011). Connexin and pannexin channel proteins are very different at the sequence level and they are not thought to be homologous, but they share similarities at the level of membrane topology (Phelan et al. 1998; Yen et al. 2007).

Invertebrates lack connexins (Phelan et al. 1998), but they have innexins which belong to the same superfamily as the pannexin proteins (Yan et al. 2007). Unlike vertebrates where gap junction and functional hemichannel capabilities have been partitioned between connexins and pannexins, innexins can function as gap junctions or in their undocked form as non-junctional membrane channels called innexons (Dahl and Muller 2014; Linden et al. 2019). Thus, innexin proteins potentially mediate both electrical and nonelectrical/chemical communication pathways and are critical for many processes including embryonic development, reproduction, and neural function (Güiza et al. 2018).

Gap junctional hemichannels are formed by connexin proteins (connexons) in vertebrates and innexin proteins (innexons) in both vertebrates and invertebrates. These two protein families are very different at the level of primary structure, but have similar topologies with four transmembrane domains (Bruzzone et al. 1996; Phelan and Starich 2001). The number of subunits per gap junctional hemichannel is variable (Oshima et al. 2016). In addition, subunit composition of gap junctional hemichannels can be homomeric (composed of a single innexin type) or heteromeric (composed of multiple innexin types). Likewise, gap junctions can be homotypic (composed of two homomeric hemichannels), heterotypic (composed of two different homomeric hemichannels), or heteromeric (composed of two heteromeric hemichannels; Figure 1C; Koval et al. 2014; Hall 2017).

Innexins are specific to animals. They are widely distributed throughout animals (Yen and Saier 2007) including ctenophores (Ryan et al. 2013; Moroz and Kohn, 2016; Slivko-Koltchik et al. 2019; Welzel and Schuster, 2022), but have not yet been identified in the genomes of sponges (Leys 2015), placozoans (Senatore et al. 2017), echinoderms (Slivko-Koltchik et al. 2019), hemichordates (Welzel and and Schuster, 2022), anthozoan cnidarians (Satterlie, 2015), and scyphozoan cnidarians (Satterlie, 2015). Phylogenetic analyses of innexins show a striking pattern of lineage-specific radiations throughout animal history (Yen and Saier 2007). Consistent with this finding, a subset of innexins from the ctenophore *Pleurobrachia bachei* were analyzed phylogenetically and shown to be more closely related to each other than to non-ctenophore innexins (Slivko-Koltchik et al. 2016). Another study identified a number of innexins in a range of ctenophore species, but none of these were complete sets from sequenced genomes and trees that included these sequences were not reported in the study (Welzel and Schuster, 2022). As such, the lack of a comprehensive phylogenetic analysis that incorporates complete genomic data from multiple ctenophore species represents a substantial gap in our understanding of innexin evolution.

While electrophysiological properties of gap junction and/or innexin hemichannels in ctenophores are unknown, high similarity of functional domains of ctenophore innexins and their relatively well studied bilaterian counterparts suggests similar physiological properties. Several basic properties of bilaterian gap junctions have been established. For example, it is known that innexin channels are nonselective, exhibit high conductance and sometimes multiple subconductance states, and are activated by intracellular calcium at physiologically relevant concentration ranges (Locovei et al. 2006; Dahl and Muller 2014).

Ultrastructural studies have offered a partial map of the anatomical distribution of gap junctions and therefore have provided insights into the functions of gap junctions in ctenophores. In *Pleurobrachia bachei*, gap junctions connect individual cilia within a comb plate (paddle-shaped bundles of thousands of cilia that comprise a comb row) suggesting a role in coordinating activity within individual plates (Satterlie and Case 1978). Satterlie and Case (1978) also observed gap junctions in the meridional canals (endoderm) that underlie the comb rows of *P. bachei*. Anctil (1985) later showed that the light-producing photocytes within the meridional canals of *Mnemiopsis leidyi* are linked to each other via gap junctions suggesting a possible role in the coordination of conduction of flashes along these canals. There is also ultrastructural evidence of gap junctions between muscle cells in *Beroe ovata* (Hernandez-Nicaise and Amsellem 1980) and *Mnemiopsis leidyi* (Hernandez-Nicaise et al. 1984). Together these data suggest that gap junctions play a key role in intercellular communication between a wide array of ctenophore cell types including neurons, photocytes, and muscle cells, and likely play a role in electrical synapses in ctenophores (Horridge 1974; Satterlie and Case 1978; Tamm 1982; Tamm 1984; Hernandez-Nicaise et al. 1989).

The current understanding of gap junctions in ctenophores is limited to ultrastructural studies and to phylogenetic analyses of a subset of innexins in a single species. Here, we phylogenetically annotate the innexins of four species of ctenophores, establish the classification and nomenclature of ctenophore innexins, show a conserved genomic architecture associated with innexin gene regulation and genomic evolution, conduct whole-mount in situ hybridization to observe the spatial expression of the four genomically clustered innexins in *Mnemiopsis*, use transcriptomic evidence to associate innexin families with specific cell types, and provide electrophysiological evidence of innexin activity. The work provides a high resolution view of the evolution of innexins in ctenophores and has implications for the evolution of animal neuromuscular systems.

## MATERIALS AND METHODS

### Transparency and Reproducibility

Prior to conducting all phylogenetic analyses, we constructed and published a phylotocol (DeBiasse and Ryan 2019) on GitHub that outlined our planned analyses. Any adjustments to the phylotocol during the course of the study have been outlined and justified in the current version of the phylotocol. These adjustments are available along with alignments, trees, and commands used in this study at: https://github.com/josephryan/ctenophore_innexins (referred to as the project GitHub repo throughout; repo snapshot available as Supplemental File 1).

### Identification of ctenophore innexins

To identify innexins in the ctenophore species *Beroe ovata, Mnemiopsis leidyi, Pleurobrachia bachei*, and *Hormiphora californensis*, we BLASTed the corresponding ctenophore protein models with Inx2 from *Drosophila melanogaster* (GenBank accession=NP_001162684.1). Any ctenophore sequence that BLASTed back to a *D. melanogaster* innexin was considered for downstream analyses. This query sequence was selected because of its relatively limited genetic change relative to other bilaterian innexins as evident from its short branch length in a recent phylogenetic study (Table SX of Abascal and Zardoya 2012). We used BLASTP with default settings and E-value cutoff of 1e-3 to search against protein models from *M. leidyi, P. bachei, H. californensis*, and B*. ovata*. The *B. ovata* innexins were recovered from an unpublished genome assembly. We also analyzed a transcriptomic dataset of *H. californensis* (prior to the publication of the *H. californensis* genome). We used TBLASTN rather than BLASTP for the transcriptomic dataset with the same E-value cutoff (1e-3). As transcriptome assemblies often contain multiple isoforms for a single gene (sometimes but not always labeled as isoforms), we generated a preliminary maximum-likelihood tree with IQ-TREE using default parameters and removed *H. californensis* innexin transcripts that had zero-length branches relative to another sister *H. californensis* innexin transcript.

### Phylogeny

We aligned each putative innexin that we identified to the Innexin PFAM domain (PF00876) and removed any sequence flanking the domain. We then aligned these sequences with MAFFT v.7.407 (Katoh and Toh 2008) using default parameters. We used this alignment to generate a maximum-likelihood tree with IQ-TREE with default parameters. We also used RAxML v.8.2.11 to generate a maximum-likelihood tree, choosing the tree with the highest likelihood value from 50 runs including 25 with starting parsimony trees and 25 with random starting trees. Lastly, we generated a Bayesian tree using MrBayes v3.26 (Ronquist and Huelsenbeck 2003). We used RAxML to generate likelihood scores for the IQ-TREE and MrBayes phylogenies and compared all four of these independent analyses, choosing the one with the highest likelihood value as our main tree. Our justification for applying multiple likelihood methodologies with multiple starting trees is that empirical analyses have shown that performance of likelihood methods and parameters are variable between datasets (Zhou et al. 2018).

The publication of the *H. californensis* genome followed the completion of our initial extensive phylogenetic analysis. To incorporate these new data into this study, we conducted a more streamlined phylogenetic analysis. As in our original analysis we aligned each putative innexin to the Innexin PFAM domain and removed flanking sequences. In cases where there were more than 1 isoform, we kept one representative based on maximizing the number of residues recognized by the PFAM domain search. Unlike in the original analysis, we did not align with MAFFT, instead we removed insertions from the alignment generated by the hmmsearch tool (with -A parameter) from HMMer (Finn et al. 2011). This approach greatly sped up the process and did not change the results of reanalyzed datasets. In this analysis we expanded the outgroups to include all the innexins from the genomes of the following species: *Branchiostoma lanceolatum* (Chordata), *Capitella teleta* (Annelida), *Lottia gigantea* (Mollusca), *Nematostella vectensis* (Cnidaria), and *Schistosoma mansoni* (Platyhelminthes). These additional sequences were all downloaded from release 51 of Ensembl Metazoa (Kinsella et al. 2011). We generated a maximum-likelihood tree from the resulting alignment using RAxML v.8.2.12 with the LG model.

### Beroe ovata genomic data

The *B. ovata* contig that contains INXB–D is from a preliminary assembly of the *B. ovata* genome and has been uploaded to the GitHub repository associated with this study. We have made the latest *B. ovata* genome assembly available at BovaDB (http://ryanlab.whitney.ufl.edu/bovadb). In addition, the *B. ovata* gene models for each of the innexins have been uploaded to this GitHub repository as well.

### Single-cell/embryo Innexin RNA expression

We used the *Mnemiopsis* Genome Project Portal (Moreland et al. 2020) to gather temporal expression information for each *M. leidyi* innexin. These temporal expression profiles were based on single-embryo developmental expression data reported in Levin et al. (2016) and Hernandez and Ryan (2018). We report the occurrence of *M. leidyi* innexins in expression clusters (approximate cell types) from adult single cell RNA-Seq data (Sebe-Pedros et al. 2018). We created a Perl script (print_coexp_all.pl in the project GitHub repo) to parse unique molecular identifier (UMI) counts from supplemental files of Sebe-Pedros et al. (2018) and count co-expression of innexins in individual cells based on these data.

### Animal culture for whole-mount in situ hybridization

*Mnemiopsis leidyi* adults were collected from floating docks in marinas surrounding the St. Augustine FL, USA area. Wild-caught animals spawned overnight in accordance with their circadian rhythm (Sasson and Ryan 2016) and their embryos were collected and reared to cydippid stages in filtered natural seawater. Cydippids were fed rotifers (*Brachionus plicatilis*, L-type, Reed Mariculture, Campbell, CA) ad libitum until they reached spawning size (~0.5-2 mm diameter). Prior to fixation, animals were starved for at least 24 hours in UV-sterilized, 1 μm-filtered natural sea water.

### RNA probe design and synthesis for whole-mount in situ hybridization

Probe templates were synthesized in vitro (by GenScript) based on known full-length coding sequences (1-1.2 kb) for each of the four target *M. leidyi* innexin genes. Probe sequences were checked against one another using Clustal Omega and with BLAST searches against the whole *M. leidyi* genome (Ryan et al. 2013) to ensure low likelihood of nonspecific binding. Digoxigenin-labeled antisense RNA probes were synthesized using the Ambion MEGAscript Kit (AM1334). Probe sequences are provided in Supplementary Table 2.

### Fixation and whole-mount in situ hybridization

Cydippids were fixed following Mitchell et al. (2021). Briefly, animals were fixed in 16% Rain-X Original Formula for 1 hour at room temperature and subsequently post-fixed with ctenophore in situ fixation buffer 1 (4% paraformaldehyde + 0.02% glutaraldehyde in FSW) for 5 minutes and ctenophore fixation buffer 2 (4% paraformaldehyde in FSW) for 1 h at room temperature in flat-bottomed, 24-well polystyrene plates. Fixed samples were dehydrated into methanol and then stored in 100% methanol at −20°C for at least 16 hours. Whole-mount in situ hybridization was performed following Pang and Martindale (2008), with detection being modified slightly using a 4:1 ratio of nitro-blue tetrazolium chloride: 5-bromo- 4-chloro-3-indolylphosphate toluidine salt. Each of the four probes were developed for at least 24 hours with the no probe control being developed as long as the slowest probe and washed with 50mM EDTA to stop the reaction. Samples were then washed several times with PTw (PBS + 0.1% (v/v) Tween-20) and then cleared in 80% glycerol at 4°C for several days. Cleared samples were imaged on a Zeiss AxioImager M2 microscope.

### *Tissue RNA*-Seq

We leveraged previously published tissue-specific RNA-Seq data from *M. leidyi* tentacle bulbs and comb rows that were reported in Babonis et al. (2018). To this we added transcriptome data from *M. leidyi* aboral organs that were collected and sequenced in the same way (all with 3 replicates) and at the same time as the other tissue data. We dipped medium-sized (20-35mm) *M. leidyi* adult individuals from floating docks in marinas surrounding the St. Augustine FL, USA area. Aboral organs were carefully excised and were snap-frozen using dry ice. RNA extraction, library preparation, and sequencing were performed by the Interdisciplinary Center for Biotechnology Research at the University of Florida. Three independent replicates, each of a single extraction from a single individual, were sequenced on a single lane of a HiSeq 3000 using a paired-end protocol. Raw sequence data have been deposited in the European Nucleotide Archive (accession PRJNA787267).

We used the rsem-calculate-expression script from RSEM version 1.3.0 (Li and Dewey, 2011) with the --bowtie2 option to align reads to the ML2.2. gene models. We used DESeq2 v1.20.0 (Love et al. 2014) to generate normalized counts in the form of transcripts per million (TPM) from these alignments.

### Electrophysiology and data analysis

We performed whole-cell voltage clamp recordings to detect and characterize the activity of putative gap junction channels in isolated *M. leidyi* muscle cells. To isolate muscle cells, we dissected small sections from adult ctenophore lobes containing muscle and mesoglea (extracellular matrix). Samples (from 15 individuals in total) were triturated with micropipette in 400-500 μL modified artificial sea water (extracellular solution: 486 mM NaCl, 5 mM KCl, 13.6 mM CaCl_2_, 9.8 mM MgCl2, 10 mM HEPES, and pH adjusted to 7.8 with NaOH or Tris-base) to separate individual cells. Dissociated cells were then plated on 35-mm Petri dishes filled with artificial sea water and allowed to settle for at least one hour before recording. We used Axiovert 100 (Carl Zeiss Inc., Germany) or Olympus IX-71 (Olympus Corp., Japan) inverted microscopes to visualize cells. Smooth muscle cells were identified morphologically by their elongated shape and numerous processes and by their ability to contract either spontaneously or after being stimulated with glutamate (Supplementary Movie 1) or high potassium solution. Intracellular solution used in whole cell recordings contained (in mM): 210 mM KCl, 696 mM Glucose, 0 mM Ca^2+^, 1mM EGTA, 10mM HEPES, with NaOH or Tris-base to adjust pH to 7.8 (intracellular low calcium solution) or 210 mM KCl, 696 mM Glucose, 0 mM EGTA, 0.001 mM CaCl_2_, 10 mM HEPES, and NaOH or Tris-base to adjust pH to 7.8 (intracellular high calcium solution). Calcium concentration in intracellular high calcium solution was likely higher than 1μM.

We pulled patch electrodes from borosilicate capillary glass (BF150-86-10, Sutter Instruments, USA) using a Flaming-Brown micropipette puller (P-87, Sutter Instruments, USA). Resistance of the electrodes was 1–3 MΩ as measured in artificial sea water. Currents were measured with either an Axopatch 200A or 200B patch-clamp amplifier (Molecular Devices, USA) using an AD–DA converter (Digidata 1320A, Molecular devices, USA), low-pass filtered at 5 kHz, and sampled at 5–20 kHz. Data were collected and analyzed with pCLAMP 9.2-10 software (Molecular Devices, USA) in combination with SigmaPlot 10-14 (SPSS, USA). Only cells characterized by a cell-attached patch seal resistance ≥1 GΩ and a relatively high input resistance (≥300 MΩ, 1.1±0.3 GΩ on average) were chosen for analysis. After establishing the whole-cell voltage clamp mode, we monitored the activity of currents at a holding potential of −60 to −70 mV for 2–5 minutes. Then the muscle cells were initially hyperpolarized by −40 to −50 mV and 200 ms voltage steps were applied in 10 mV increments. All current traces free from the activity of voltage-gated channels, typically in the range −120-(−40) mV, were carefully reviewed on possible high conductance channel activity. We performed all recordings at room temperature.

We used single channel currents to generate current-voltage relationships. The reversal potential estimates based on single channel currents are more accurate since the direct measurement of unitary currents eliminates or reduces the necessity for pharmacological dissection of integral currents (Bobkov et al., 2011). The reversal potential estimates for potassium (V_r_,K^+^), chloride (V_r_,Cl^−^) and monovalent cation (V_r_,X^+^) selective channels were calculated using Nernst equation.

## RESULTS

### Innexins in ctenophores

We identified 9 innexins in *Pleurobrachia bachei*, 19 in *Hormiphora californensis*, and 12 in both *Mnemiopsis leidyi* and *Beroe ovata.* We performed a range of maximum-likelihood and Bayesian analyses to phylogenetically classify these innexins. We found that all ctenophore innexins are most closely related to other ctenophore innexins (Figure 2A) and have therefore radiated in parallel to the innexins of other animals (Figure 2B). The ctenophore innexins have substantial conservation in the four regions predicted to be the transmembrane helices, contain the four highly conserved extracellular cysteines, and include the conserved proline in the second transmembrane domain (Supplementary Figure 1).

**Figure 2.**
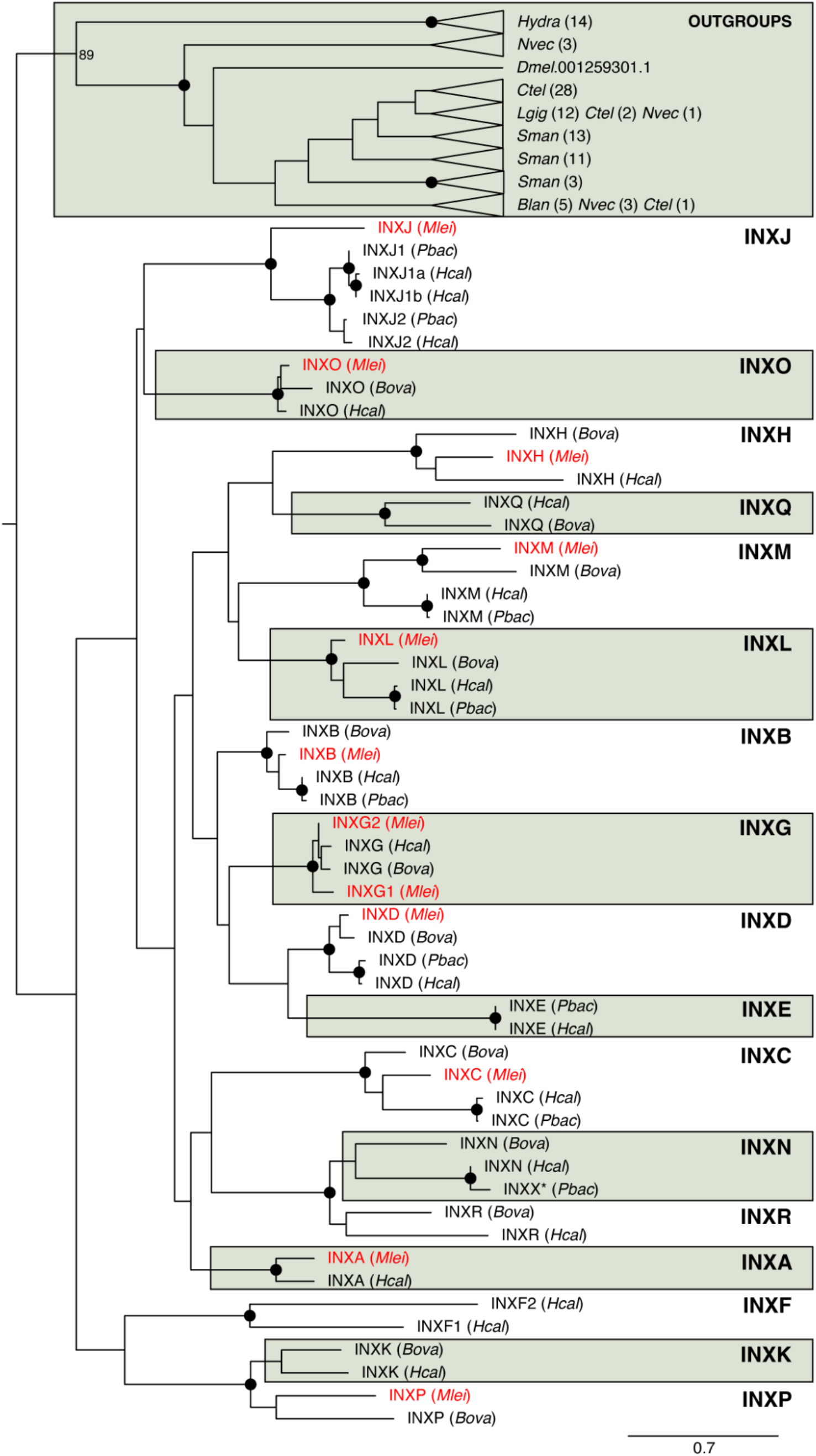
Evolution of ctenophore innexins. (A) Maximum likelihood tree of innexins from four ctenophore species *Mnemiopsis leidyi* (Ml), *Pleurobrachia bachei* (Pb), *Beroe ovata* (Bo), and *Hormiphora californensis* (Hc) as well as full sets of innexins from non-ctenophores including *Hydra vulgaris* (Cnidaria), *Nematostella vectensis* (Cnidaria), *Capitella teleta* (Annelida), *Lottia gigantea* (Mollusca), *Schistosoma mansoni* (Platyhelminthes), *Branchiostoma lanceolatum* (Chordata), as well as Inx2 from *Drosophila melanogaster* (Arthropoda). Solid circles at the nodes indicate bootstrap support greater than or equal to 90%. A version of this tree with all bootstrap values and without collapsed outgroup clades is available as Supplementary Figure 2.

Based on our phylogeny, we identified 17 ctenophore innexin families that we have named INXA–INXR (we did not include an INXI to avoid confusion with INX1 in other species; Figure 2A). To infer gene gains/losses (Figure 2B), we assumed that absence from a clade indicates a historical gene loss and multiple genes from the same species in a single clade indicates a historical gene duplication (parsimony principles). We infer that 14 innexin families (i.e., INXA–D,G,H,J–O,Q,R) arose in the stem ancestor of Ctenophora, INXP arose in the last common ancestor of *B. ovata* and *M. leidyi*, INXE arose in the last common ancestor of *P. bachei* and *H. californensis*, and INXF arose in the *H. californensis* lineage. We inferred 13 gene losses within these ctenophore lineages including four in the *M. leidyi* lineage (INXK, N, Q, and R), two in the *B. ovata* lineage (INXA and J), seven in the lineage leading to *P. bachei* (INXA, G, H, K, O, Q, and R) and none in the lineage leading to *H. californensis*. We identified four duplications including one in the lineage leading to *P. bachei* and *H. californensis*, one in the *M. leidyi* lineage (INXG), and two in the *H. californensis* lineage (INXJ and F). Names and accessions of ctenophore innexins are provided in Supplementary Table 3.

### Conserved genomic cluster of Innexins

The INXB, INXC, and INXD genes are within 40 kilobases of each other in each of the *M. leidyi, P. bachei*, and *B. ovata* genomes (Figure 3A). These clusters include no detectable intervening non-innexin genes. In *M. leidyi*, a fourth innexin, INXA, is less than 40 Kb upstream of the INXB-D cluster. Between INXA and INXB are three non-innexins, an ankyrin-related gene (ML25994a), an undescribed ctenophore specific gene (ML25995a), and an intraflagellar transport-related gene (ML25996a). The *B. ovata* INXB-D cluster spans 33 Kb. INXA was not recovered in *B. ovata* or *P. bachei*.

**Figure 3:**
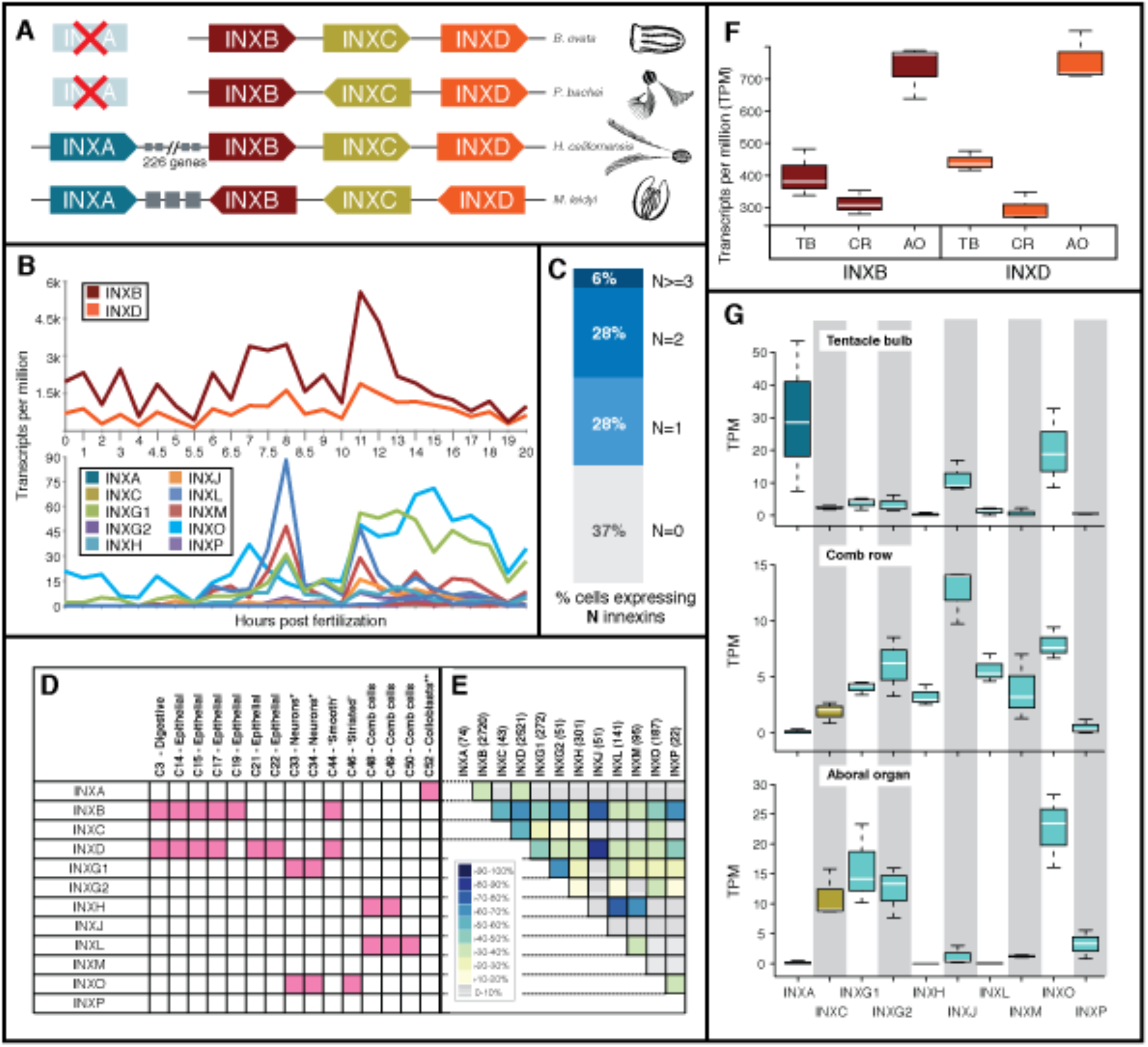
temporal, cellular, and spatial expression of Innexins. (A) Innexin clusters in the genomes of *Beroe ovata, Pleurobrachia bachei, Hormiphora californensis*, and *Mnemiopsis leidyi*. *Mnemiopsis leidyi* has a four-gene Innexin cluster that includes INXA, INXB, INXC, and INXD. The genomes of *B. ovata* and *P. bachei* have a cluster that includes INXB, INXC, and INXD. Grey boxes represent 3 non-innexin genes between INXA and INXB in *M. leidyi*. All four of these innexins are on chromosome 10 in *H. californensis*, but there are 226 genes separating INXA and INXB. The clusters are not to scale. In all genomes INXB, INXC, and INXD are within 20Kb of each other and in *M.leidyi*, the entire cluster including INXA is less than 80Kb. (B) Temporal gene expression in single *M. leidyi* embryos during the first 20 hours of development (Levin et al. 2016) shows that INXB and INXD are both highly expressed and are tightly coordinated (top section). Comb plates are formed at 8 hours post fertilization and at this point there is a spike in expression of several innexin genes that are expressed in comb-plate cell types and tissues (i.e., INXL, INXM, INXG.1, INXH, INXJ, and INXP). (C) Percentage of single cells expressing 0, 1, 2, or >3 innexins (Supplementary Table 4). (D) Columns represent metacells from single-cell data (Sebe-Pedros et al. 2018). Pink squares represent expression of the corresponding innexin in 50% or more of the cells that make up the specified metacell (full counts per metacell are in Supplementary Table 5). (E) Heatmap of co-expression of each innexin in individual cells. The number of individual cells that express each innexin are in parentheses (in the column header). The percentage was determined by taking the number of cells with co-expression divided by the lowest number when comparing the number of individual cells that express each innexin (full co-expression counts are in Supplementary Table 6). (F) Expression of *M. leidyi* INXB and INXD in three replicates of bulk tissue RNA-Seq from tentacle bulbs (TB), comb rows (CR), and aboral organ (AO). INXB and INXD are shown separately because they are very highly expressed relative to the other innexins. (G) Expression of *M. leidyi* innexins in three replicates of bulk tissue RNA-Seq from tentacle bulbs, comb rows, and aboral organ. * Metacells C33 and C34 were hypothesized to be neurons in Sachkova et al. (2021). ** Metacells C52 were hypothesized to be colloblasts in Babonis et al. (2018).

INXB, INXC, and INXD are adjacent on chromosome 10 of the chromosome-level assembly of the recently sequenced genome of *Hormiphora californensis* (Schultz et al. 2021). There are 226 genes between INXA and the INXB-D cluster in *H. californensis*. Microsynteny is rare between *H. californensis* and *M. leidyi* with the largest identifiable blocks of gene microsynteny including only four genes (Schultz et al. 2021), but the block that includes INXB, INXC, and INXD is conserved and also includes a fourth gene (ML259910a in *M. leidyi* and Hcv1.av93.c10.g249.i1 in *H. californensis*), which is similar to the FAM166B gene in humans. Interestingly, INXN (Hcv1.av93.c10.g18.i1), which was lost in *M. leidyi*, is next to INXA in *H. californensis* and INXG (Hcv1.av93.c10.g192.i1) is situated in between these two sets of innexins (173 genes downstream from INXA and 54 genes upstream of INXB) suggesting the cluster was once more extensive.

### Innexin gene expression

We compared the gene expression profiles of *M. leidyi* innexins during early development, in adult tissues, and in adult single cells. In all of these data, INXB and INXD expression levels are orders of magnitude higher than all other innexins (Figure 3B). Our single-embryo RNA-Seq time-course data (Levin et al. 2016; Hernandez and Ryan 2018) show that INXB and INXD have highly coordinated expression patterns throughout development (Figure 3B).

We identified evidence of innexin co-expression in the single-cell RNA-Seq data in *M. leidyi* from Sebé-Pedrós et al. (2018). In this study, the gene expression profiles of individual cells from a dissociated adult animal were clustered to construct metacells, which approximate cell types. Any gene expressed in at least 50% of the cells comprising a metacell were considered marker genes. INXB and INXD were considered co-marker genes in a digestive (C3), ‘smooth’ muscle (C44), and 3 epithelial metacells (C14,15,17; Figure 3D). INXO and INXG1 were considered marker genes in two neural metacells (C33,34) INXH and INXL, were considered marker genes for two comb plate metacells (C48,49; Figure 3D; Supplementary Table 4).

Beyond metacells, we analyzed innexin expression in individual cells using the UMI data from the supplemental data of Sebé-Pedrós et al. (2018). The majority of the cells in this dataset express at least one innexin (63%), 34% of cells express at least two innexins, and one cell expresses 9 innexins (Figure 3C; Supplementary Table 5). These numbers are likely an underestimate as the depth of sequencing in this study (average of 36,000 reads per cell and 5 reads per UMI) was low compared to more recent single-cell RNA-Seq studies. We identified 10 instances where 50% of the cells expressing a specific innexin also expressed another innexin. Most of these pairings involve the highly expressed INXB and INXD genes including: INXB-INXC, INXB-INXG2, INXB-INXP, INXB-INXJ, INXD-INXJ, INXD-INXC, but several pairings did not include INXB and INXD including: INXG1-INXG2, INXH-INXA, INXH-INXL, INXH-INXM (Figure 3E; Supplementary Table 6).

Like the developmental time course data, the single-cell RNA-Seq data lend support to the hypothesis that the innexin genome cluster (Figure 3A) has a role in coordinating expression of INXA–D. For example, INXB is expressed in 1,761 (70%) of the cells that express INXD (Figure 3E; Supplementary Table 6). In addition, INXB and INXD are expressed in 53.5% of cells expressing INXC and in 37.8% of cells expressing INXA (Supplementary Table 6). It is important to note that while INXC is not a defining gene for any metacells (likely due to more moderate levels of expression compared to INXB and INXD) and therefore lacks any shaded cells in Figure 3D, it is expressed widely (Figure 4F).

**Figure 4.**
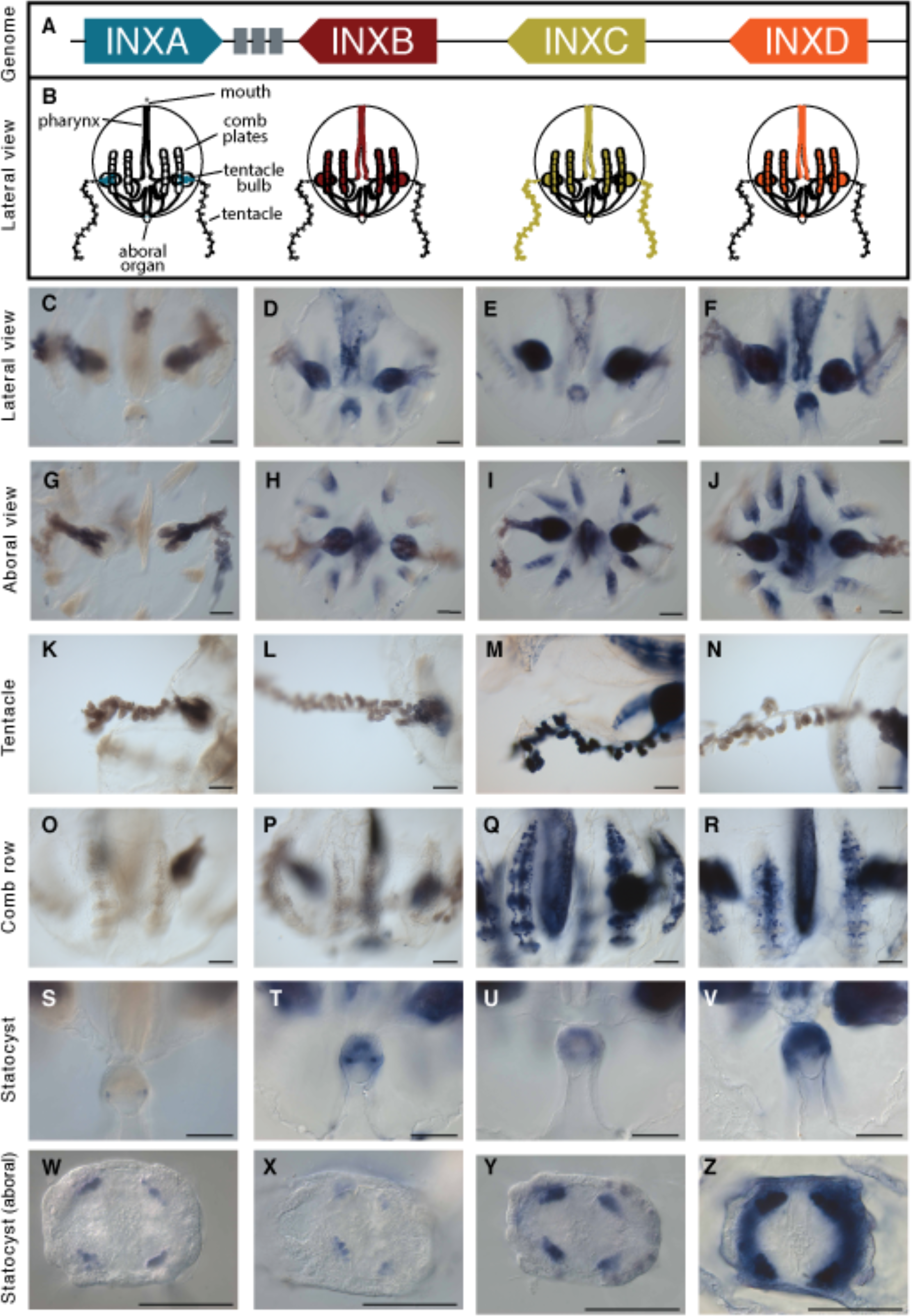
Whole-mount in situ hybridization for four clustered *Mnemiopsis leidyi* innexin genes. (A) Cartoon depiction of the innexin genomic cluster. (B) Cartoon representation of spatial patterns of expression of INXA-D. Specific patterns within expression domains of INXB-D are not shown due to slight variations in patterns between individuals. Seemingly overlapping domains in cartoons may not indicate co-expression in the same cells. (C-Z) In situ expression of INXA-D. The label to the left of each row describes the view or tissue under focus. Columns correspond to positions of genes in the genomic cluster of panel A. (C,G,K) INXA is highly expressed in the lateral ridge of the tentacle bulb. (O) INXA is not expressed in the comb rows or underlying canals. (S,W) There are four distinct INXA domains of expression in the aboral organ. (D,H) INXB is widely expressed in the tentacle bulbs, pharynx, aboral organ, and meridional canals underlying the longitudinal comb rows. (L) INXB is expressed in the tentacle bulb, but not the tentacle. (P) There is punctate INXB expression in the meridional canals underlying the comb rows. (T,X) There are four distinct INXA domains of expression in the aboral organ. (E,I) INXC is widely expressed in the tentacle bulbs, comb rows, pharynx, and aboral organ. (M) INXC has a distinct expression domain in the tentacles. (Q) INXC is expressed in the comb rows and in the underlying meridional canals. (U, Y) There are four distinct INXA domains of expression in the aboral organ. (F, J) INXD is widely expressed in the tentacle bulbs, comb rows, pharynx, and aboral organ. (N) INXD is not expressed in the tentacles. (R) There is punctate INXD expression in the meridional canals underlying the comb rows. (V, Z) INXD is expressed in a ring in the aboral organ. Coloration in the tentacles of INXA, INXB, and INXD is likely background. No probe controls are in Supplementary Figures 4–5. Patterns are representative of replicates (Supplementary Figures 6–9). All scale bars represent 100 μm.

We next identified co-expression of *M. leidyi* innexins in tissue-specific RNA-Seq data from tentacle bulbs and comb rows (Babonis et al 2018) as well as aboral organs (this study; principal components analysis in Supplementary Figure 3). As in our other expression data, INXB and INXD are both highly expressed relative to other innexins and are expressed at similar relative levels in each tissue (i.e. both are expressed higher in aboral organ tissue than in tentacle bulbs and lowest in comb rows; Figure 3 F–G). The tissue RNA-Seq expression patterns also bolster evidence for other innexins that might be working together in either a heteromeric or heterotypic capacity. For example, INXL and INXH, which were implicated in the single-cell RNA-Seq data as being involved in comb plate cells, are expressed at relatively high levels in comb row tissues and at very low levels or not at all in aboral organ and tentacle bulb samples (Figure 3G).

### Spatial expression of clustered innexin genes

We examined the localization of mRNA expression of INXA–D by whole-mount in situ hybridization in cydippid-stage *Mnemiopsis leidyi* (Figure 4). INXA expression, which is present only in one of the colloblast cell types in the single-cell data and is highest in the tentacle bulbs in our tissue RNA-Seq, is present almost exclusively in the tentacle bulbs in our in situ expression analyses. In particular, the expression is localized to the lateral ridge of the tentacle bulb and the tentacle root (Figure 4G, 4K). This expression domain in combination with the single-cell data is consistent with INXA being expressed in developing colloblasts. INXC on the other hand is expressed in a wider domain in the tentacle bulbs (Figure 4E, 4I) and is also clearly expressed in the tentacles themselves (Figure 4M). INXC is not expressed in colloblasts in the single cell data, but is most highly expressed in single cells hypothesized to be neurons by Sachkova et al. (2021; Supplementary Table 5). Together these data suggest INXC may be being expressed in neurons of the tentacles (single-cell RNA sequencing at a greater depth of sequencing is needed to confirm). In addition to the tentacle bulb expression, INXA is expressed in a small number of cells making up four distinct domains in the floor of the aboral organ (Figure 4S, 4W). INXC is also expressed in cells comprising 4 domains in the floor of the aboral organ (Figure 4G, 4W).

INXB and INXD are expressed in the tentacle bulbs, comb rows, pharynx, and aboral organ (Figure 3 and 4). Like INXA and C, INXB is expressed in a small number of cells comprising four distinct domains in the floor of the aboral organ (Figure 4T, 4X). INXD aboral expression appears to form a ring in the aboral organ (Figure 4V, 4Z). INXB-D all exhibit a speckled expression pattern in the pharynx (Figure 4D-F) and in the comb row region (Figure 4P-R).

### Electrophysiological evidence of innexin function in Mnemiopsis leidyi

To begin to functionally characterize the ctenophore innexins, we tested whether these channels, like their bilaterian counterparts (pannexins, innexins), are capable of forming functional non-junctional channels (innexons). We tested whether the conductance of channels in *M. leidyi* muscle cells was consistent with the single channel-conductance parameters estimated for the innexon channels of other species (250-550pS, in 150mM K+, Locovei et al. 2006; Bao et al. 2007; Kienitz et al. 2011). We leveraged the high conductance of innexon channels to visualize activity of single innexons (channel unitary currents) and characterize *the* channel gating.

We used isolated muscle cells because these are abundant in primary cell cultures from oral lobe tissue and are easily identifiable both morphologically and functionally. These cells contract upon excitation (Supplementary Movie 1), express innexins (Sebe-Pedros et al. 2018; Figure 5) as well as voltage-gated channels typical for excitable cells (Sebe-Pedros et al. 2018; Figure 5A,C; Supplementary Figure10) and are sensitive to amino acids such as glutamate/glycine (Alberstein et al. 2015; Supplementary Figure 11 and Supplementary Movie 1).

**Figure 5.**
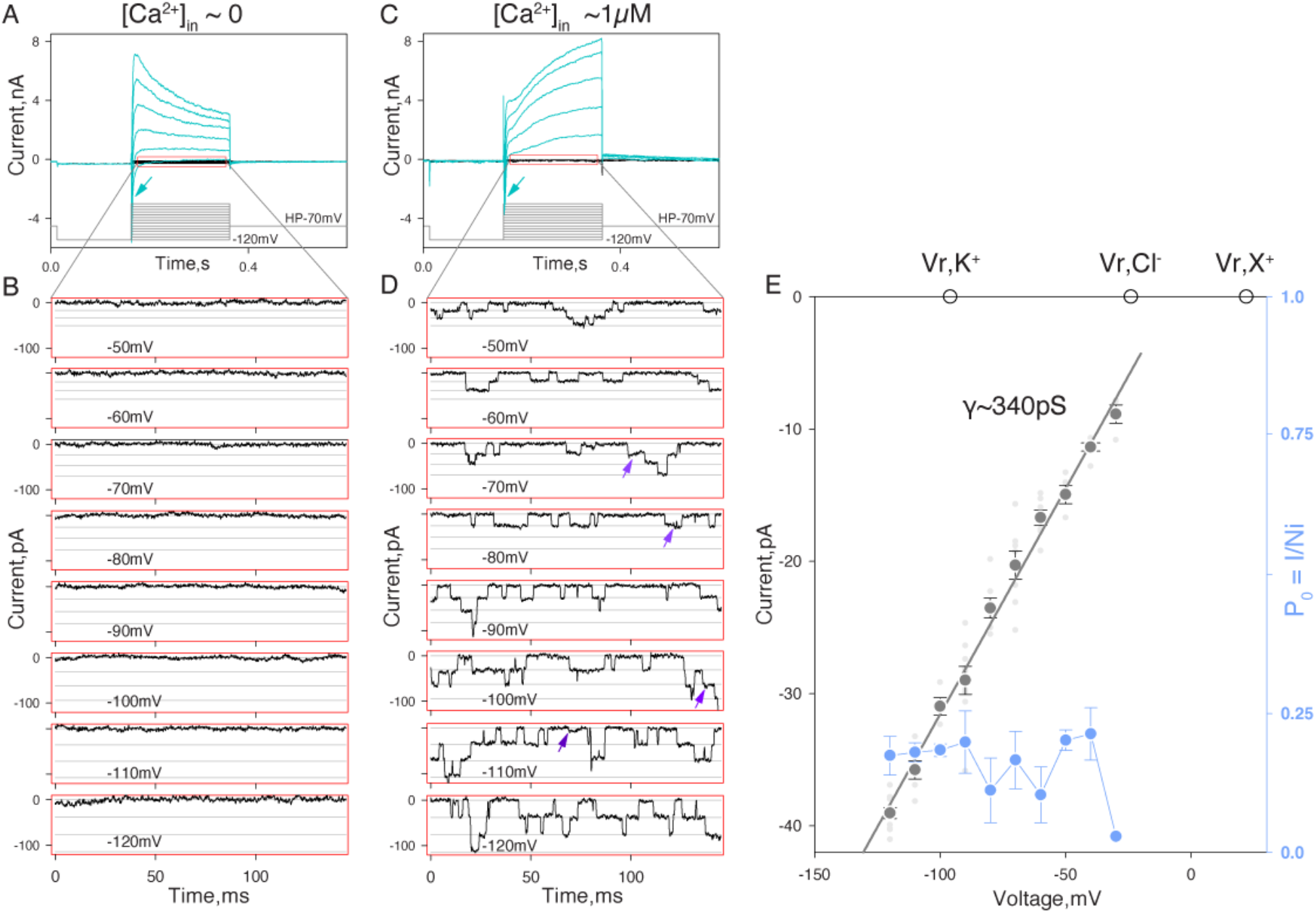
Activity of putative innexons in *Mnemiopsis leidyi* muscle cells. Representative whole-cell currents recorded from isolated muscle cells in low (A,B) and high (C,D) intracellular calcium. (A,C) Voltage protocol diagram showing muscle cells were initially hyperpolarized by −50mV and then 200ms voltage steps were applied in 10mV increments. Voltage gated currents are depicted in cyan (also Supplementary Figure 10 for details). For example, inward currents characterized by fast activation/inactivation kinetics (arrows) represent activity of voltage-gated sodium channels. Innexin channel activity outlined in red represents traces without active voltage gated channels which are shown in more detail in B and D. (B,D) Each panel displays portions of current traces obtained at different potentials (as indicated). Horizontal lines depict unitary current levels. Arrows depict possible short-lived subconductance states. Current values (y-axis) represent current values minus the basal current level. (E) Plot of the relationship between current (pA) and voltage (mV) based on the mean values (dark grey symbols) of single channel current amplitudes obtained at different voltages (as in panel D). Data were obtained from 10 cells in total. Each light grey symbol represents a single channel amplitude estimated for an individual cell. A linear approximation of this relationship corresponds to a slope conductance of ~340 pS. Empty circles depict the predicted reversal potentials of ideal potassium (V_r_,K^+^), chloride (V_r_,Cl^−^) and monovalent cation (V_r_,X^+^) selective channels in the given experimental conditions. Note, unitary current-voltage relationship suggests nonselective nature of the channel pore. Providing estimates of the relative permeability of the channel for inorganic and organic ions would require further detailed analysis. The blue symbols and lines (right y-axis) in panel D represent voltage dependence of channel open probability expressed as P_o_=I/Ni (where I = integral current, N = number of channels detected in given conditions, i = single channel amplitude). Voltage dependence of P_o_ was analyzed for 4 cells except for −30mV where n=1. Extracellular conditions in A and B are identical. Data presented were filtered at 1 kHz and reduced 10-fold.

We used whole-cell voltage clamp mode to record whole-cell currents and detect potential activity of innexons. We focused exclusively on recording innexon unitary currents. To better resolve innexon unitary currents in the whole-cell mode, only cells characterized by relatively high input resistance (≥300MOhm) and cell-attached patch seal resistance ≥1GOhm were chosen for analysis.

We tested for potential innexon channel activity in 0 intracellular Ca^2+^_free_ (1mM EGTA +0 Ca^2+^ added) condition using the experimental paradigm described in methods. In 9 out of 9 cells chosen for analysis we found no indication of high conductance channel activity (Fig5.A, B). These results suggest that *M. leidyi* innexon channels require intracellular calcium for activation (e.g., Locovei et al. 2006; Bao et al. 2007).

We therefore tested whether these channels can be activated by relatively high intracellular calcium (≥1μM). In 10 out of 24 cells examined, we were able to reliably detect channel activity characterized by parameters potentially matching the following basic properties of bilaterian innexin hemichannels: high single-channel conductance (~340pS; Fig 5B-C), the apparent lack of ion selectivity (inferred reversal potential near 0; Fig 5C), the potential sensitivity to intracellular calcium, and the apparent lack of voltage-dependent channel gating between −120 and −30mV (Fig 5E, blue line and symbols, right Y-scale).

## DISCUSSION

Our phylogenetic analyses suggest that innexins independently radiated within Ctenophora. This pattern of lineage-specific diversification within major animal lineages is an evolutionary tendency of innexins (Yen and Saier 2007) as it has also been described in insects (Hasegawa and Turnbull 2014), tunicates, nematodes (Suzuki et al. 2010), molluscs, annelids, and cnidarians (Fushiki et al. 2010; Abascal and Zardoya, 2013). This pattern of independent diversification in multiple animal lineages is unusual. On the contrary, most gene superfamilies consist of a combination of early-established families (i.e., families that arose in the stem ancestors of ancient lineages like Metazoa, Parahoxozoa, and Bilateria) and more recently established families (i.e., those that arose from lineage-specific expansions). Examples of this more common pattern of expansion can be seen in homeoboxes (Ryan et al. 2006), Wnts (Pang et al. 2011), LIM genes (Koch et al. 2012), trypsins (Babonis et al. 2019), opsins (Schnitzler et al. 2012), voltage-gated ion channels (Moran et al. 2015), and most other large superfamilies. It is possible that this pattern of diversification in innexins is the byproduct of processes that led to the establishment of major animal lineages. Alternatively, given that lineage-specific gene family expansion is often associated with adaptation and biological innovation (Lespinet et al 2002), it is possible that the diversification of innexins in the stem lineages of many major animal clades played a fundamental role in establishing these lineages

From analyses of public *M. leidyi* single-cell RNA-Seq data, we show that most cells express at least one innexin and more than 35% of cells express more than one innexin. This is similar to what is seen in other animals (e.g., *Caenorhabditis elegans* Altun et al. 2009; *Hydra vulgaris* Seibert et al. 2019) and suggests innexin channels have played a role in many cell types since the last common animal ancestor.

The identification of a cluster of innexins in the genomes of *M. leidyi, P. bachei* and *B. ovata* is a rare instance of such a conserved cluster of genes in ctenophores. As such, these data offer an example of how an extensive gene family arose via tandem duplication in ctenophores. In addition, when combined with expression data, the cluster provides a foundational example of genomic structure providing a functional role (i.e., gene regulation) in ctenophores. The expression data suggest that genes within the cluster are co-regulated in *M. leidyi*. INXB, INXC, and INXD have highly overlapping spatial patterns and are often expressed within the same cells. There is also single-cell evidence for overlapping expression between INXA and INXB, as well as INXA and INXD. These overlapping expression profiles suggest that there is regulatory architecture under purifying selection that is maintaining this cluster throughout long periods of evolutionary time. It is curious that the INXC expression overlaps with INXB and INXD, but that in all of the quantifiable expression experiments (i.e., single-embryo, single-cell, and tissue RNA-Seq) INXC is expressed at a much lower level than INXB and INXD.

The combined expression profiles of the highly expressed INXB and INXD suggest that these genes are expressed in many cells, often in the same cell, and that overall they are expressed in a consistent 2 to 1 ratio. The INXC gene expression profile spatially overlaps with INXB and INXD, but the combined data point to this gene being expressed at much lower levels than INXB and INXD. These expression data combined with the extraordinary conserved nature of this gene cluster within ctenophores, suggests a shared regulatory mechanism and presents the best such foothold from which to interrogate gene regulation from within ctenophores. We hypothesize that the coordinated expression of these innexin genes has some bearing on the subunit makeup of the resulting channels and potential gap junctions. Given that the cluster has been maintained over millions of years, we further hypothesize that a similar regulatory system and subunit makeup was present in the last common ancestor of all ctenophores (at least in the last common ancestor of the four ctenophores we analyzed).

Our in situ gene expression data suggests that innexins are expressed in adjacent domains of the aboral organ (Figure 4W–Z). It is difficult to discern from these data whether the comb row expression of INXB–D involves comb plate cilia, gametes, photocytes or other cell types located in this region. In published single-cell RNA-Seq data (Sebe-Pedros et al. 2018), INXB and INXD are considered marker genes for clusters of cells (metacells) labeled as epithelial, digestive, and muscle cells (Figure 3D), but INXB and INXD are also highly expressed in many other cells including several metacells that were not labeled in the original study (Figure 3E).

While significant progress has been made characterizing the functional, electrophysiological properties of gap junction channels in bilaterians (e.g. Wang et al. 2014; Bhattacharya et al. 2019; Walker and Schafer 2020; for review Dahl and Muller 2014; Skerrett and Williams 2017 and Guiza et al. 2018) relatively little is known about gap junction channels in non-bilaterian lineages. Here we provide the first demonstration of the activity of putative innexon channels from ctenophore muscle cells. The smooth muscle cells of *M. leidyi* are excellent cells to investigate innexins: there is published ultrastructural evidence of gap junctions connecting these cells (Hernandez-Nicaise et al. 1984), most smooth muscles sampled in published RNA-Seq data express INXB (87 of 186 cells) and INXD (78 of 186 cells), and the electrophysiological and molecular properties can be studied in great detail due to the ability to maintain these cells in primary cell culture (Supplementary Figure 10).

Potential effects of intracellular calcium ions on gap junctional channels are likely more complex than were initially suggested (e.g., Deleze and Loewenstein 1976; Loewenstein and Rose 1978; reviewed by Skerrett and Williams 2017) and not limited to suppression of electrical and/or dye coupling by elevated cytoplasmic calcium. Indeed, activation by intracellular calcium in physiologically relevant concentration range appears to be an important common property of non-junctional pannexins and innexins (Locovei et al. 2006; Bao et al. 2007; Kienitz et al. 2011; Dahl and Muller 2014).

Our results outline the following basic physiological properties of *M. leidyi* innexin channels: (1) high single-channel conductance, (2) the presence of subconductance states, (3) the apparent lack of ion selectivity, (4) the potential sensitivity to intracellular calcium, and (5) the apparent lack of voltage dependency of channel gating (at least between −120 to −30 mV).

While individually, these properties could be attributed to other channel types, collectively, they are consistent with properties of gap junction channels. For example, Maxi-Cl channels (SLCO2A1, a member of the solute carrier organic anion transporter family) are expressed in ctenophore muscle cells (e.g. ML18358a is expressed in 22% of a metacell identified as muscle–c43–in Sebe-Pedros et al. 2018) and would also be characterized by high unitary conductance, the occurrence of subconductance states, and calcium dependent activity, but these channels are anion-selective and exhibit a distinct voltage dependence of their open probability (Sabirov et al. 2017). Similarly, we can eliminate the potential involvement of large conductance calcium-activated potassium channels (ML128229a is expressed in 47% of a muscle metacell–c47) because of the ion selectivity of these channels. We can also eliminate inositol 1,4,5-trisphosphate receptor (insP3R) channels (ML25824a shows no expression in muscle metacells above background) since these are calcium sensitive, nonselective cation channels, require cytoplasmic IP3 to be active, and otherwise show extremely low basal activity (Dellis et al. 2006). Furthermore, the relatively high concentration of divalent cations used in our experiments would dramatically decrease unitary conductance in InsP3R channels (Bezprozvanny and Ehrlich 1994; Mak and Foskett 1998). Thus we conclude that the properties of the channels we characterize here, collectively, are consistent with those of innexons.

To more rigorously implicate the innexins in the ctenophore intercellular signaling, future efforts will require exploring the detailed functional properties and pharmacological and molecular profiles of signaling pathways involved. Our electrophysiological results will have a substantial impact on orienting future efforts to uncover the role of innexins in distributing signals throughout ctenophore neuro-muscular and neuro-sensory networks.

## CONCLUSION

Our data show that *M. leidyi* innexins are expressed widely and some at very high levels in almost every cell type. This is consistent with ultrastructural studies (Satterlie and Case, 1978; Hernandez-Nicaise and Amsellem, 1980; Hernandez-Nicaise et al. 1984; Anctil, 1985) showing that, like in other animals (e.g. Hall, 2017), gap junctions are broadly deployed throughout the ctenophore body plan. Our whole-cell recordings of *M. leidyi* smooth muscle cells show channel activity consistent with the channel activity of gap junction channels in bilaterians. The genomic clustering of innexins suggests an ancient regulatory mechanism underlying innexin expression. Together these data support a key role for innexins and gap junctions in the biology of ctenophores and provide an essential starting point for future exploration of innexins, genome regulation, and gap junctions in ctenophores.

## ACKNOWLEDGEMENTS

We thank Michel Anctil for helpful conversations. We thank David Ortiz for help generating Figure 3. We thank Barry Ache for a critical reading of parts of this manuscript. We thank James Strother for access to equipment and conversations. The views expressed in this manuscript do not necessarily reflect the views of those acknowledged.

This work was supported by the National Science Foundation (DEB-1542597 to J.F.R.) and the National Science Foundation Research Experience for Undergraduates (REU) Program (DBI-1156528 to J.F.R.). This research was supported by an Allen Distinguished Investigator Award, a Paul G. Allen Frontiers Group advised grant of the Paul G. Allen Family Foundation to J.F.R. and M.Q.M. A.E. was supported by a National Science Foundation Postdoctoral Research Fellowship in Biology (DBI-2010755). The funders had no role in study design, data collection and analysis, decision to publish, or preparation of the manuscript.

