## supplemental material for "Independent innexin radiation shaped signaling in ctenophores"

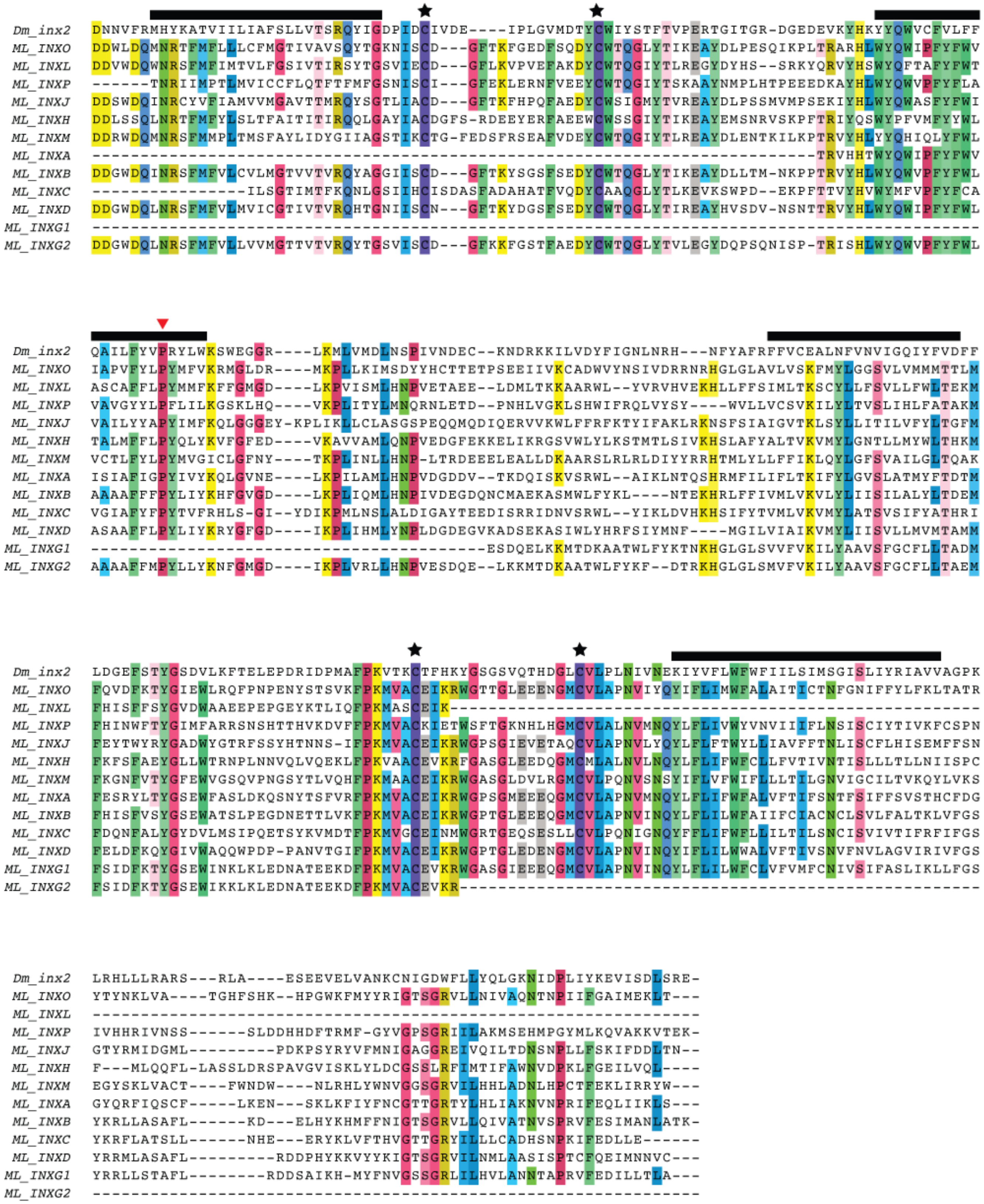
Dm_inx2 DNNVFRMHYKATVIILIAFSLLVTSRQYIGDPIDCIVDE----IPLGVMDTYCWIYSTFTVPERTGITGR-DGEDEVKYHKYYQWVCFVLFF
ML_INXO DDWLDQMNRTFMFLLLCFMGTIVAVSQYTGKNISCD---GFTKFGEDFSQDYCWTQGLYTIKEAYDLPESQIKPLTRARHLWYQWIPFYFWV
ML_INXL DDVWDQWNRSFMFIMTVLFGSIVTIRSYTGSVIECD---GFLKVPVEFAKDYCWTQGIYTLREGYDYHS--SRKYQRVYHSWYQFTAFYFWT
ML_INXP ------TNRIIMPTLMVICCFLQTFTFMFGSNISCI---GFEKLERNFVEEYCWTQGIYTSKAAYNMPLHTPEEEDKAYHLWYQWVPFYFLA
ML_INXJ DDSWDQINRCYVFIAMVVMGAVTTMRQYSGTLIACD---GFTKFHPQFAEDYCWSIGMYTVREAYDLPSSMVMPSEKIYHLWYQWASFYFWI
ML_INXH DDLSSQLNRTFMFYLSLTFAITITIRQQLGAYIACDGFS-RDEEYERFAEEWCWSSGIYTIKEAYEMSNRVSKPFTRIYQSWYPFVMFYYWL
ML_INXM DDRWDQMNRSFMMPLTMSFAYLIDYGIIAGSTIKCTG-FEDSFRSEAFVDEYCWTQGIYTLREAYDLENTKILKPTRVYHLYYQHIQLYFWL
ML_INXA ---------------------------------------------------------------------------TRVHHTWYQWIPFYFWV
ML_INXB DDGWDQINRSFMFVLCVLMGTVVTVRQYAGGIISCD---GFTKYSGSFSEDYCWTQGLYTIKEAYDLLTM-NKPPTRVYHLWYQWVPFYFWL
ML_INXC ----------------ILSGTIMTFKQNLGSIIHCISDASFADAHATFVQDYCAAQGLYTLKEVKSWPD--EKPFTTVYHVWYMFVPFYFCA
ML_INXD DDGWDQLNRSFMFVLMVICGTIVTVRQHTGNIISCN---GFTKYDGSFSEDYCWTQGLYTIREAYHVSDV-NSNTTRVYHLWYQWIPFYFWL
ML_INXG1 -------------------------------------------------------------------------------------------
ML_INXG2 DDGWDQLNRSFMFVLLVVMGTTVTVRQYTGSVISCD---GFKKFGSTFAEDYCWTQGLYTVLEGYDQPSQNISP-TRISHLWYQWVPFYFWL
Dm_inx2 QAILFYVPRYLWKSWEGGR----LKMLVMDLNSPIVNDEC--KNDRKKILVDYFIGNLNRH---NFYAFRFFVCEALNFVNVIGQIYFVDFF
ML_INXO IAPVFYLPYMFVKRMGLDR----MKPLLKIMSDYYHCTTETPSEEIIVKCADWVYNSIVDRRNRHGLGLAVLVSKFMYLGGSVLVMMMTTLM
ML_INXL ASCAFFLPYMMFKFFGMGD----LKPVISMLHNPVETAEE--LDMLTKKAARWL--YVRVHVEKHLLFFSIMLTKSCYLLFSVLLFWLTEKM
ML_INXP VAVGYYLPFLILKGSKLHQ----VKPLITYLMNQRNLETD--PNHLVGKLSHWIFRQLVYSY----WVLLVCSVKILYLTVSLIHLFATAKM
ML_INXJ VAILYYAPYIMFKQLGGGEY-KPLIKLLCLASGSPEQQMQDIQERVVKWLFFRFKTYIFAKLRKNSFSIAIGVTKLSYLLITILVFYLTGFM
ML_INXH TALMFFLPYQLYKVFGFED----VKAVVAMLQNPVEDGFEKKELIKRGSVWLYLKSTMTLSIVKHSLAFYALTVKVMYLGNTLLMYWLTHKM
ML_INXM VCTLFYLPYMVGICLGFNY----TKPLINLLHNP-LTRDEEELEALLDKAARSLRLRLDIYYRRHTMLYLLFFIKLQYLGFSVAILGLTQAK
ML_INXA ISIAFIGPYIVYKQLGVNE----LKPILAMLHNPVDGDDV--TKDQISKVSRWL--AIKLNTQSHRMFILIFLTKIFYLGVSLATMYFTDTM
ML_INXB AAAAFFFPYLIYKHFGVGD----LKPLIQMLHNPIVDEGDQNCMAEKASMWLFYKL----NTEKHRLFFIVMLVKVLYLIISILALYLTDEM
ML_INXC VGIAFYFPYTVFRHLS-GI--YDIKPMLNSLALDIGAYTEEDISRRIDNVSRWL--YIKLDVHKHSIFYTVMLVKVMYLATSVSIFYATHRI
ML_INXD ASAAFFLPYLIYKRYGFGD----IKPLIHMLYNPLDGDEGVKADSEKASIWLYHRFSIYMNF----MGILVIAIKVMYLIISVLLMVMTAMM
ML_INXG1 ----------------------------------------ESDQELKKMTDKAATWLFYKTNKHGLGLSVVFVKILYAAVSFGCFLLTADM
ML_INXG2 AAAAFFMPYLLYKNFGMGD----IKPLVRLLHNPVESDQE--LKKMTDKAATWLFYKF--DTRKHGLGLSMVFVKILYAAVSFGCFLLTAEM
Dm_inx2 LDGEFSTYGSDVLKFTELEPDRIDPMAFPKVTKCTFHKYGSGSVQTHDGLCVLPLNIVNEKIYVFLWFWFIILSIMSGISLIYRIAVVAGPK
ML_INXO FQVDFKTYGIEWLRQFPNPENYSTSVKFPKMVACEIKRWGTTGLEEENGMCVLAPNVIYQYIFLIMWFALAITICTNFGNIFFYLFKLTATR
ML_INXL FHISFFSYGVDWAAEEPEPGEYKTLIQFPKMASCEIK--------------------------------------------------------
ML_INXP FHINWFTYGIMFARRSNSHTTHVKDVFFPKMVACKIETWSFTGKNHLHGMCVLALNVMNQYLFLIVWYVNVIIFLNSISCIYTIVKFCSPN
ML_INXJ FEYTWYRYGADWYGTRFSSYHTNNS-IFPKMVACEIKRWGPSGIEVETAQCVLAPNVLYQYLFLFTWYLLIAVFFTNLISCFLHISEMFFSN
ML_INXH FKFSFAEYGLLWTRNPLNNVQLVQEKLFPKVAACEVKRFGASGLEEDQGMCMLALNVLNQYLFLIFWFCLLFVTIVNTISLLLTLLNIISPC
ML_INXM FKGNFVTYGFEWVGSQVPNGSYTLVQHFPKMAACEIKRWGASGLDVLRGMCVLPQNVSNSYIFLVFWIFLLLTILGNVIGCILTVKQYLVKS
ML_INXA FESRYLTYGSEWFASLDKQSNYTSFVRFPKMVACEIKRWGPSGMEEEQGMCVLAPNVMNQYLFLIFWFALVFTIFSNTFSIFFSVSTHCFDG
ML_INXB FHISFVSYGSEWATSLPEGDNETTLVKFPKMVACEIKRWGPTGLEEEQGMCVLAPNVINQYLFLILWFAIIFCIACNCLSVLFALTKLVFGS
ML_INXC FDQNFALYGYDVLMSIPQETSYKVMDTFPKMVGCEINMWGRTGEQSESLLCVLPQNIGNQYFFLIFWFLLILTILSNCISVIVTIFRFIFGS
ML_INXD FELDFKQYGIVWAQQWPDP-PANVTGIFPKMVACEIKRWGPTGLEDENGMCVLAPNVINQYIFLILWWALVFTIVSNVFNVLAGVIRIVFGS
ML_INXG1 FSIDFKTYGSEWINKLKLEDNATEEKDFPKMVACEVKRWGASGIEEEQGMCVLAPNVINQYLFLILWFCLVFVMFCNIVSIFASLIKLLFGS
ML_INXG2 FSIDFKTYGSEWIKKLKLEDNATEEKDFPKMVACEVKR------------------------------------------------------
Dm_inx2 LRHLLLRARS---RLA----ESEEVELVANKCNIGDWFLLYQLGKNIDPLIYKEVISDLSRE-
ML_INXO YTYNKLVA----TGHFSHK--HPGWKFMYYRIGTSGRVLLNIVAQNTNPIIFGAIMEKLT---
ML_INXL ---------------------------------------------------------------
ML_INXP IVHHRIVNSS------SLDDHHDFTRMF-GYVGPSGRIILAKMSEHMPGYMLKQVAKKVTEK-
ML_INXJ GTYRMIDGML---------PDKPSYRYVFMNIGAGGREIVQILTDNSNPLLFSKIFDDLTN--
ML_INXH F---MLQQFL-LASSLDRSPAVGVISKLYLDCGSSLRFIMTIFAWNVDPKLFGEILVQL----
ML_INXM EGYSKLVACT----FWNDW----NLRHLYWNVGGSGRVILHHLADNLHPCTFEKLIRRYW---
ML_INXA GYQRFIQSCF-----LKEN---SKLKFIYFNCGTTGRTYLHLIAKNVNPRIFEQLIIKLS---
ML_INXB YKRLLASAFL------KD---ELHYKHMFFNIGTSGRVLLQIVATNVSPRVFESIMANLATK-
ML_INXC YKRFLATSLL------NHE---ERYKLVFTHVGTTGRYILLLCADHSNPKIFEDLLE------
ML_INXD YRRMLASAFL---------RDDPHYKKVYYKIGTSGRVILNMLAASISPTCFQEIMNNVC---
ML_INXG1 YRRLLSTAFL--------RDDSAIKH-MYFNVGSSGRLILHVLANNTAPRVFEDILLTLA---
ML_INXG2 ---------------------------------------------------------------

**Supplementary Figure 1. Alignment of *Drosophila melanogaster* Inx-2 and all of the *Mnemiopsis leidyi* innexins reveals conserved residues.** Conserved features of *Drosophila melanagoster* Inx2 (Dm_Inx2) are indicated above the alignment (after Tazuke et al. 2002). Extracellular cysteine residues are indicated by stars and conserved proline in the second transmembrane domain are indicated by a red triangle. The N-terminal and C-terminal ends of these sequences are not shown and were not used in the phylogenetic analyses. A background color is used for columns where 50% or more of residues are the same (default colors used in Unipro UGENE).

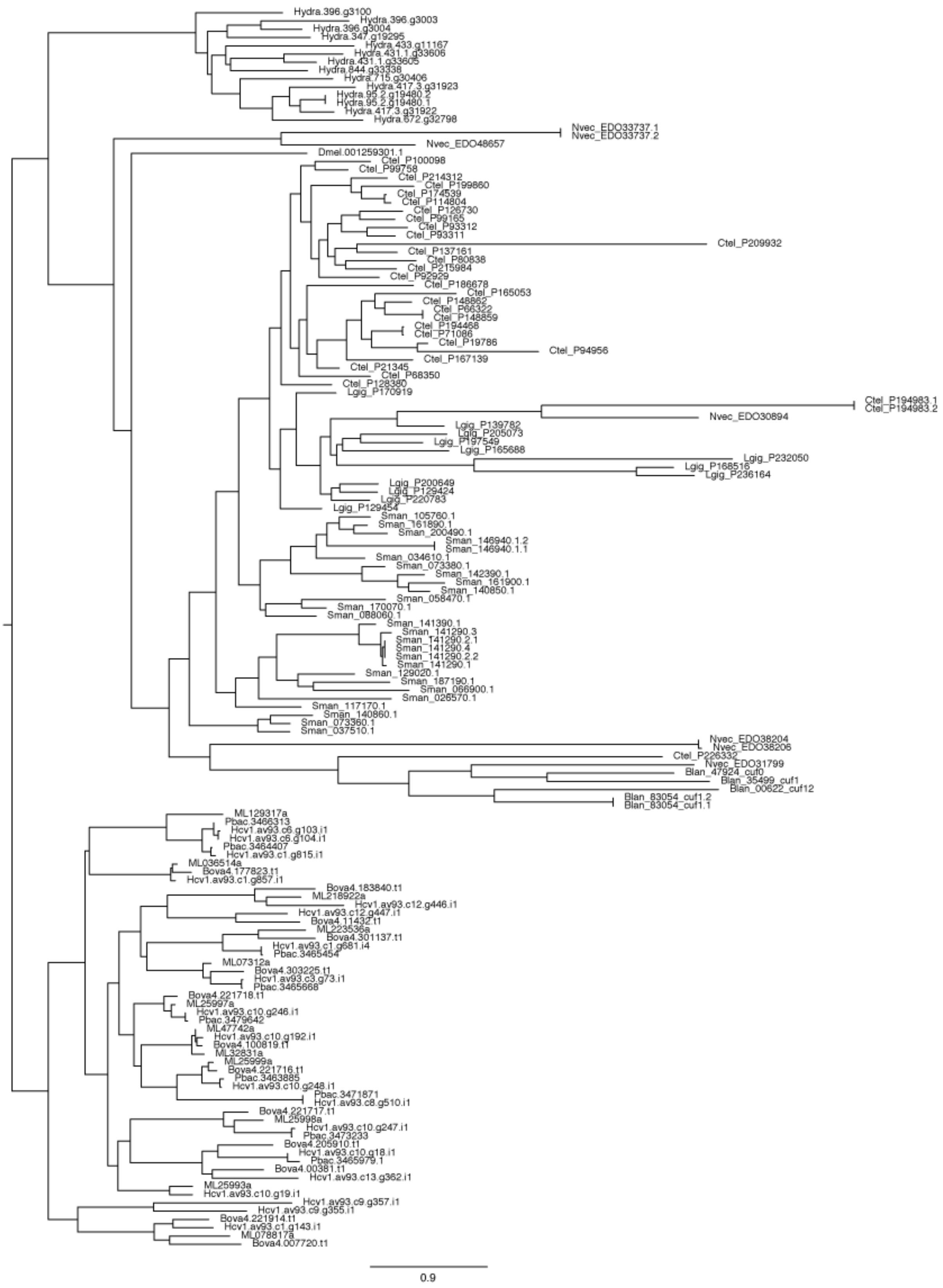


**Supplementary Figure 2. Detailed maximum-likelihood phylogeny of innexins.** This tree is summarized in Figure 2 of the main manuscript. Bootstrap values below 70 are regarded as suspect and below 50 are regarded as unreliable. It is included to show details of collapsed clades. A newick version of this tree (6taxa_plus_hcal_genome_plus_bilat.pruned.tre) is available in the Github repository associated with this study.

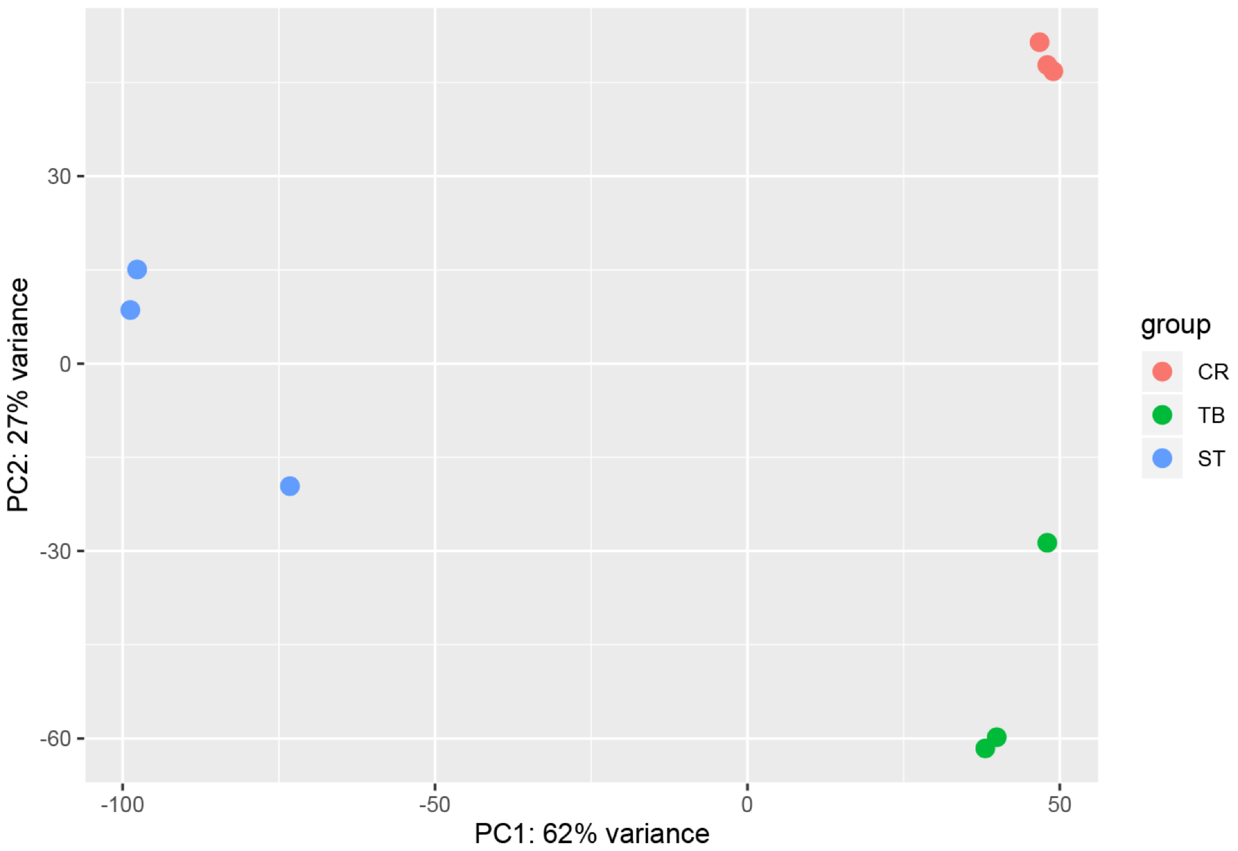


**Supplementary Figure 3. Principal components analysis of tissue RNA-Seq data.** PCA was performed using plotPCA function in DESeq2 on genes with counts in at least 3 libraries. CR=comb row, TB=tentacle bulb, ST=statocyst.

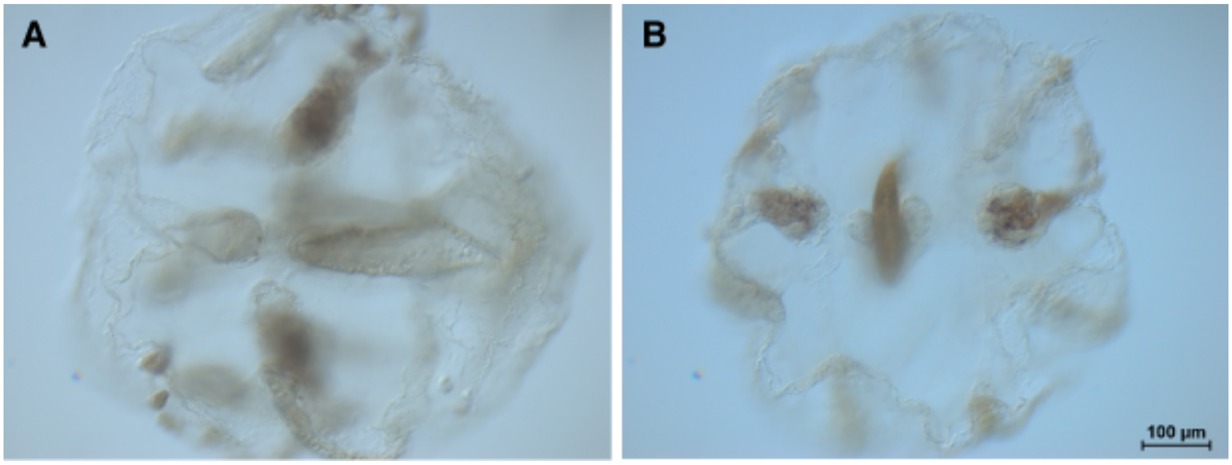


**Supplementary Figure 4. No probe controls for whole mount in situ hybridization.** Slight background staining can be seen in pharynx, tentacle, and tentacle bulb tissue. (A) lateral view, (B) aboral view.

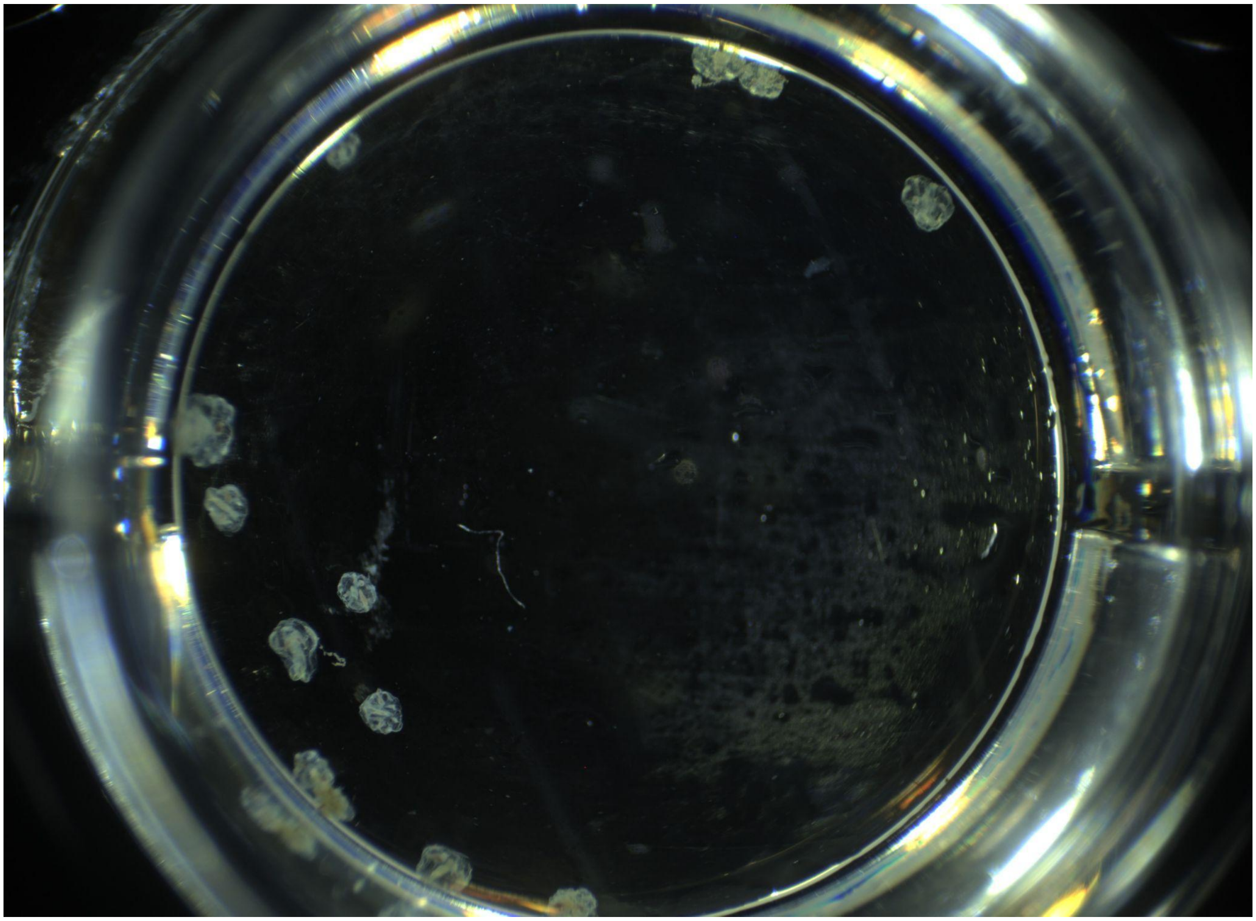

**Supplementary Figure 5. No probe controls for whole mount in situ hybridization in *M.leidyi* cydippids.** Image of a well from a 24-well tissue culture plate (well diameter=16.2mm). The slight background staining seen in Supplementary Figure 2 can be seen in most of these cydippids.

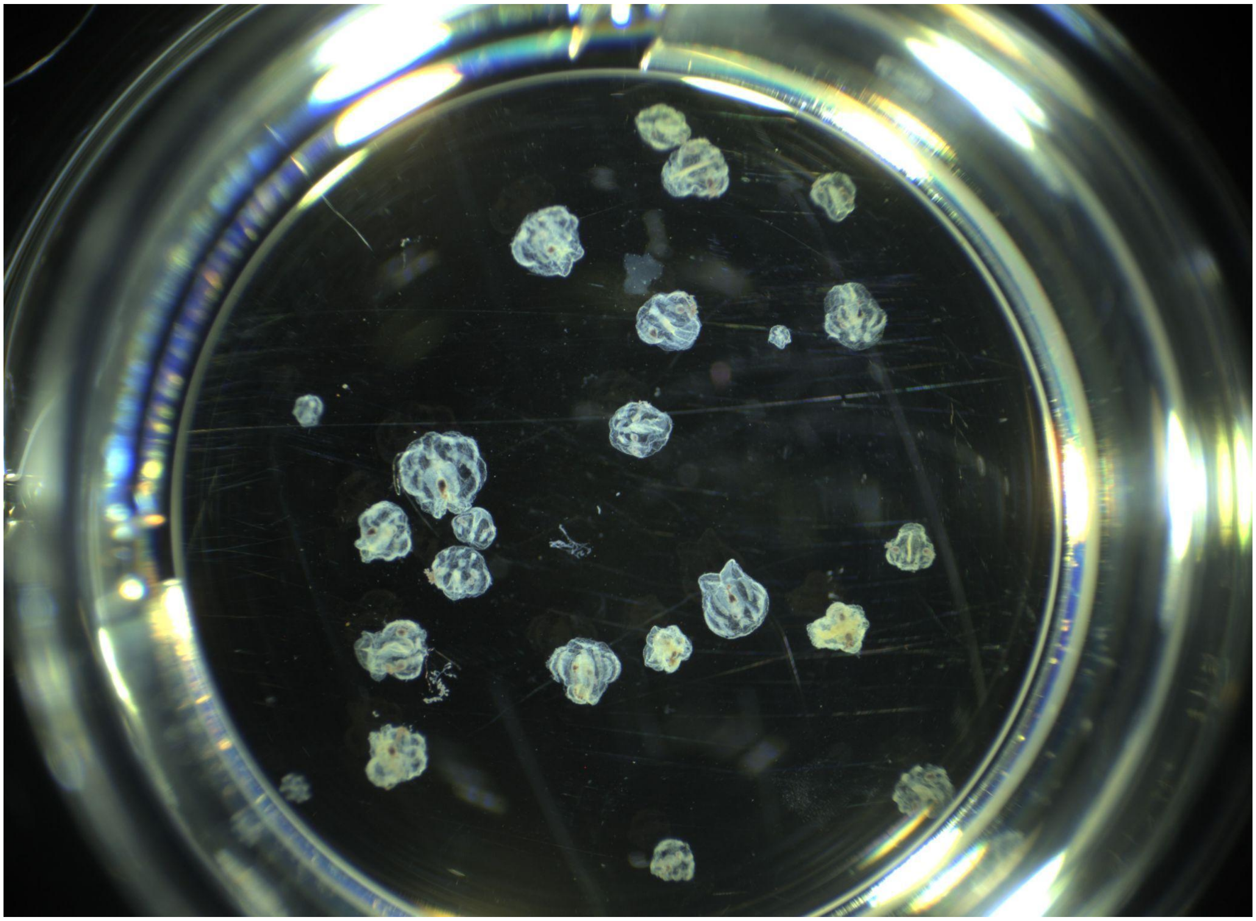

**Supplementary Figure 6. Whole mount in situ hybridization with INXA (ML25993a) probe in *M.leidyi* cydippids.** Image of a well from a 24-well tissue culture plate (well diameter=16.2mm). The in situ pattern observed in Figure 4 for INXA is representative of patterns seen in most of these individuals.

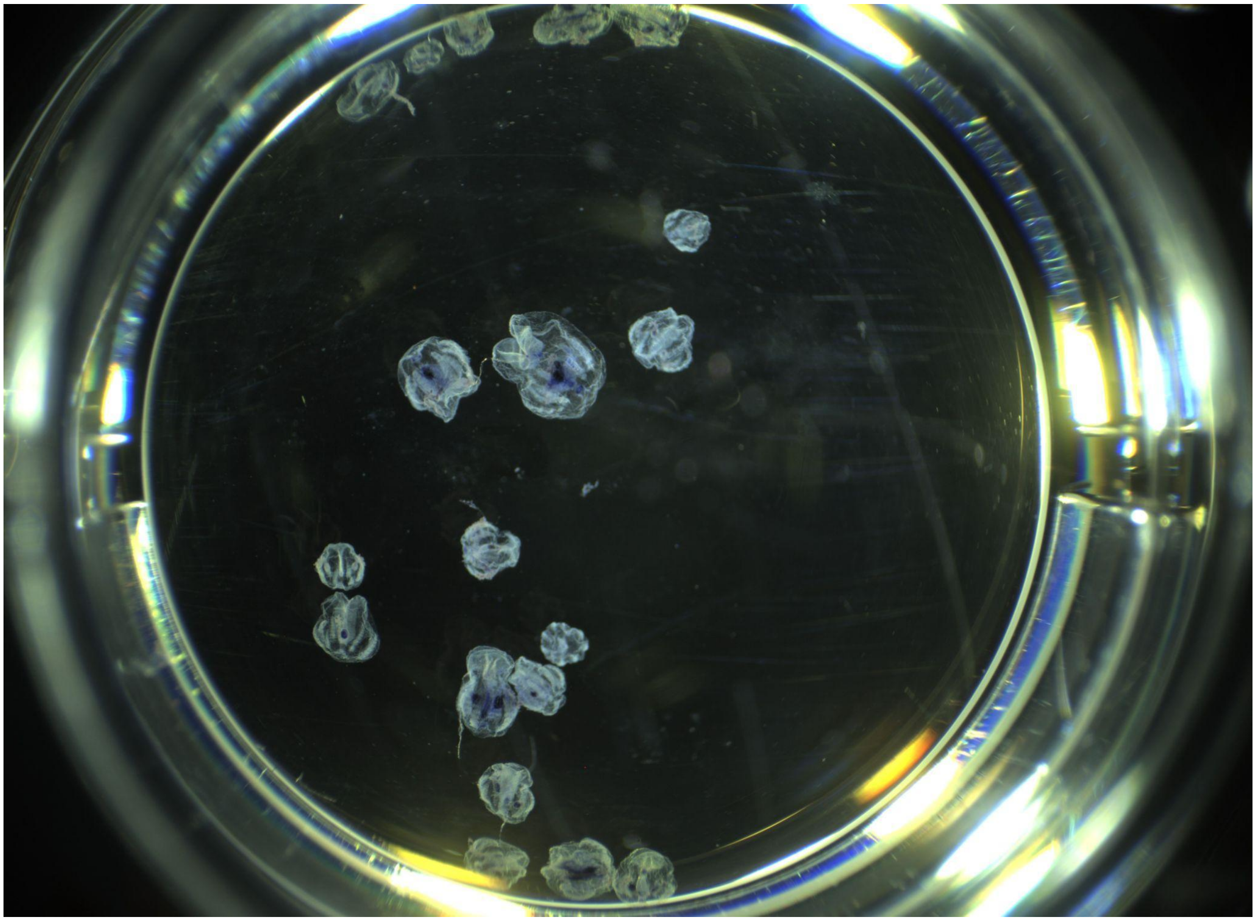

**Supplementary Figure 7. Whole mount in situ hybridization with INXB (ML25997a) probe in *M.leidyi* cydippids.** Image of a well from a 24-well tissue culture plate (well diameter=16.2mm). The in situ pattern observed in Figure 4 for INXB is representative of patterns seen in most of these cydippids.

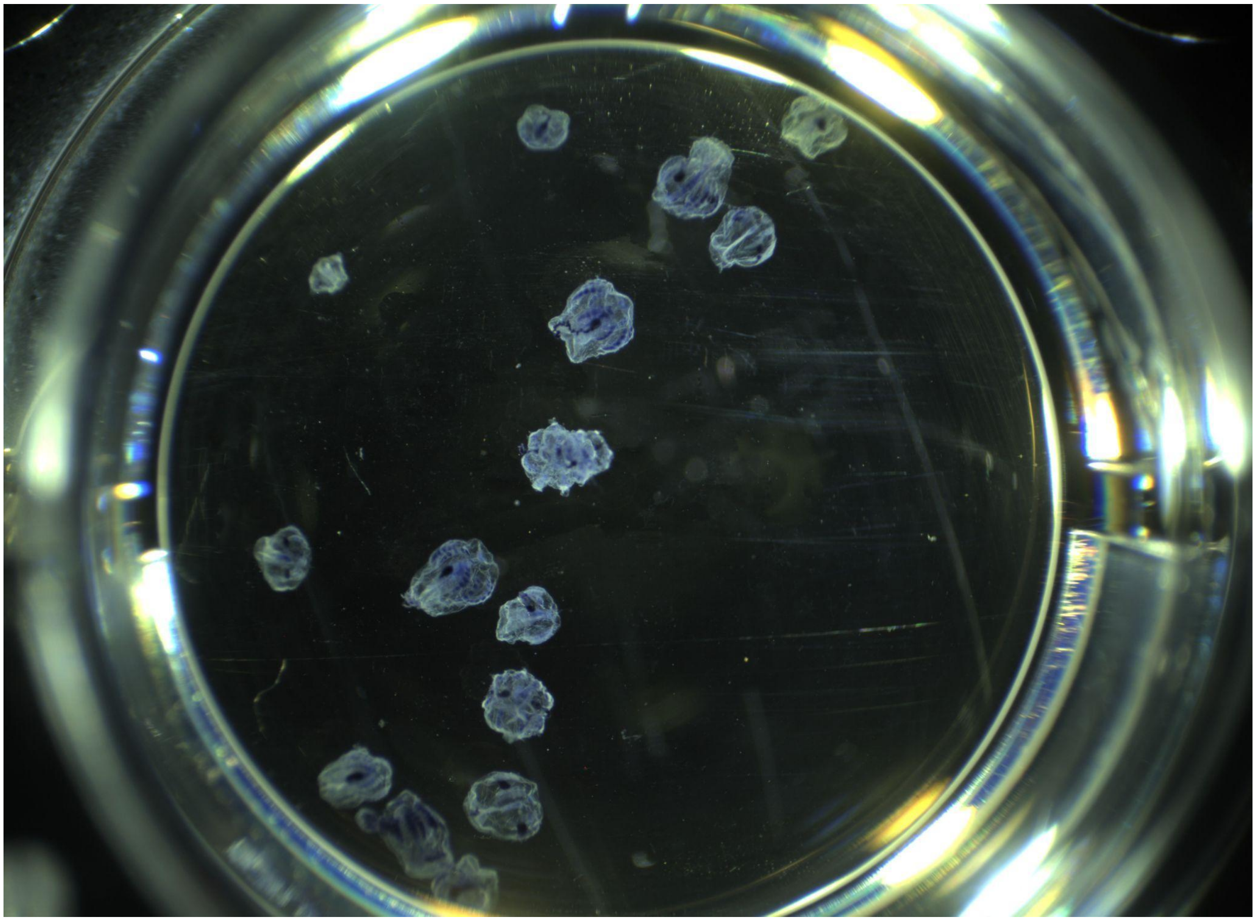

**Supplementary Figure 8. Whole mount in situ hybridization with INXC (ML25998a) probe in *M.leidyi* cydippids.** Image of a well from a 24-well tissue culture plate (well diameter=16.2mm). The in situ pattern observed in Figure 4 for INXC is representative of patterns seen in most of these cydippids.

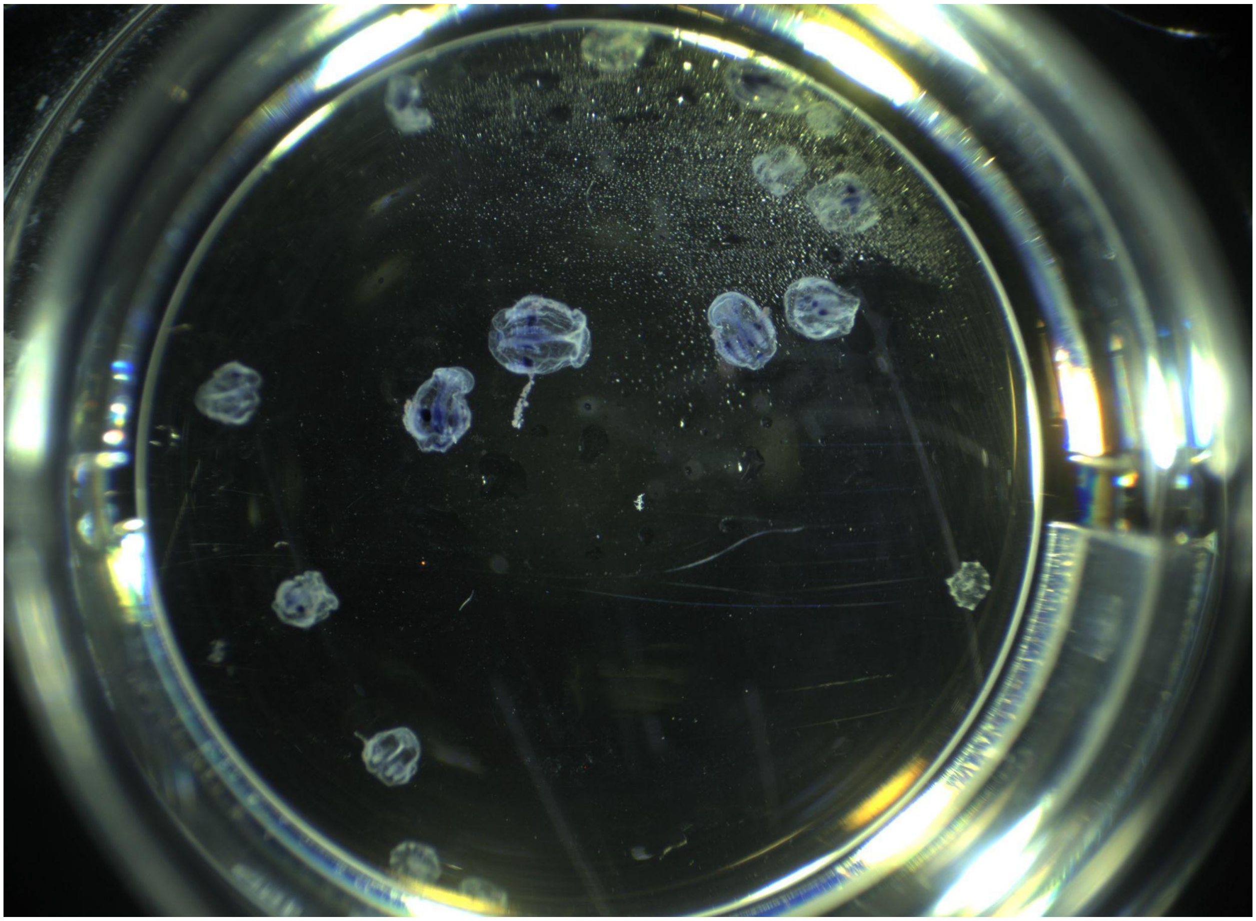

**Supplementary Figure 9. Whole mount in situ hybridization with INXD (ML25999a) probe in *M.leidyi* cydippids.** Image of a well from a 24-well tissue culture plate (well diameter=16.2mm). The in situ pattern observed in Figure 4 for INXD is representative of patterns seen in most of these cydippids.

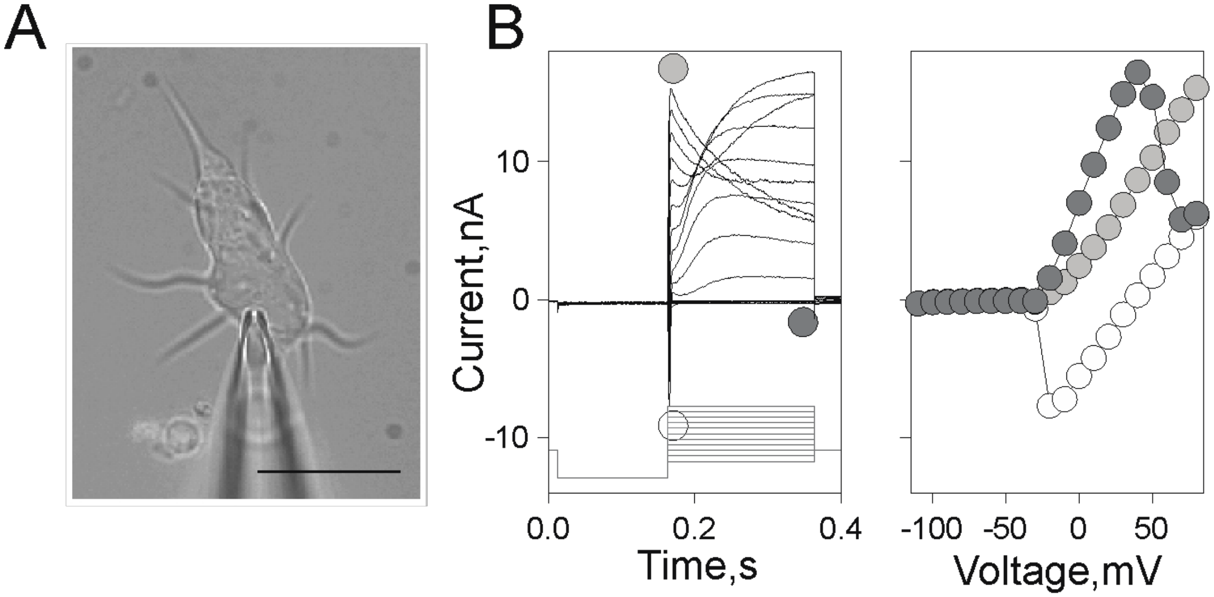


**Supplemental Figure 10. General characteristics of the ctenophore *Mnemiopsis leidyi* muscle cell conductances**. (A) Muscle cell/muscle cell fragment in the ctenophore primary cell culture. (B) Typical set of voltage-dependent currents recorded from isolated muscle cells using whole-cell voltage-clamp recording: Inward currents characterized by fast activation/inactivation kinetics (white circle/s) represent activity of voltage-gated sodium channels. Fast activating outward currents (A type currents, grey circle/s) are presumably mediated by the activity of delayed rectifier potassium channels. And, relatively slowly activating outward currents (dark grey circle/s) reflect the activity of voltage-gated, calcium sensitive potassium channels. (B, right panel) Respective current-voltage characteristics. Current scales in B are the same. Scale bar 20um. A and B represent different cells.

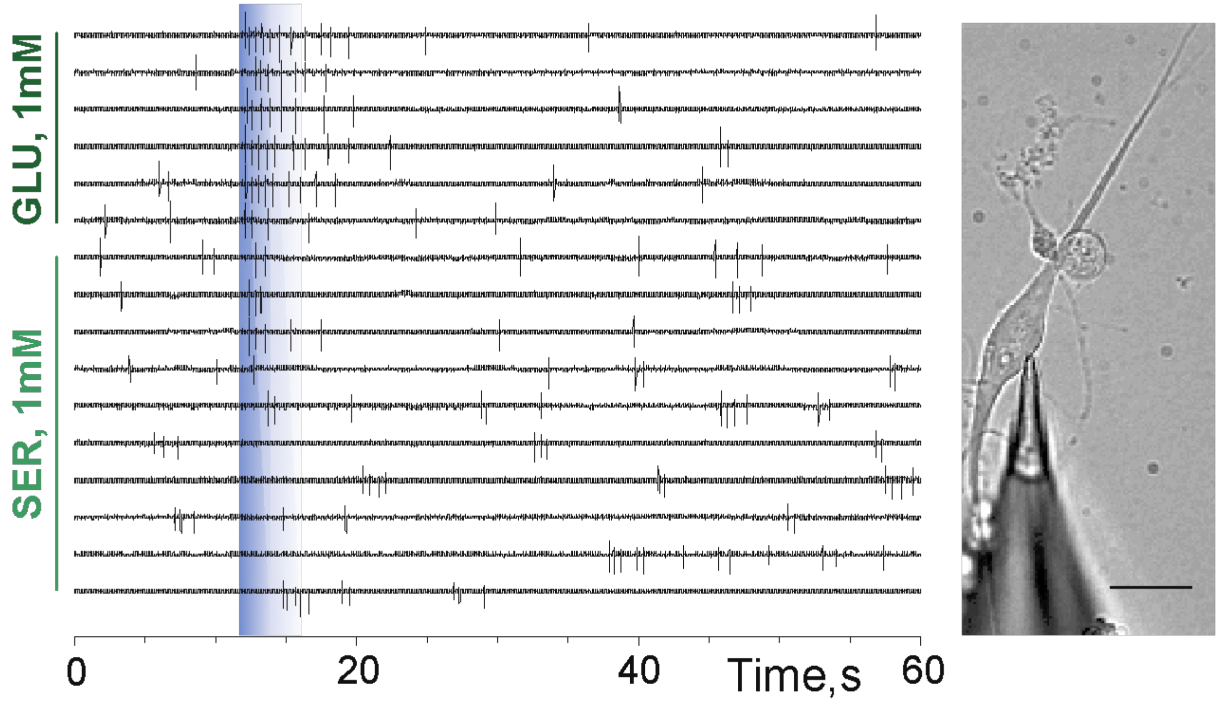


**Supplemental Figure 11. Ligand evoked activity of isolated ctenophore *Mnemiopsis leidyi* muscle cells.** Typical responses of ctenophore muscle cells to 500ms pulses of a transmitter application (blue faded bar) were recorded using loose-patch clamp configuration. These cells are sensitive to glutamate and capable of generating action potentials in a consistent manner. Glutamate repetitively increases ongoing rate of discharge. Multi-channel rapid solution changer (RSC-160, Bio-Logic) under software control (Clampex 9/10, Molecular Devices) was used for neurotransmitter/s application. Simultaneous video recording (Supplemental Video 1) shows evoked contractile activity of the cell. Scale bar 20um.

**Supplementary Table 1. Peer reviewed publications providing phylogenomic evidence supporting ctenophores as the sister group to the rest of animals**

| |
|---|
| Dunn CW, Hejnol A, Matus DQ, Pang K, Browne WE, Smith SA, Seaver E, Rouse GW, Obst M, Edgecombe GD, Sørensen MV, Haddock SH, Schmidt-Rhaesa A, Okusu A, Kristensen RM, Wheeler WC, Martindale MQ, Giribet G. Broad phylogenomic sampling improves resolution of the animal tree of life. Nature. 2008 Apr 10;452(7188):745-9. doi: 10.1038/nature06614 |
| Hejnol A, Obst M, Stamatakis A, Ott M, Rouse GW, Edgecombe GD, Martinez P, Baguñà J, Bailly X, Jondelius U, Wiens M, Müller WE, Seaver E, Wheeler WC, Martindale MQ, Giribet G, Dunn CW. Assessing the root of bilaterian animals with scalable phylogenomic methods. Proc Biol Sci. 2009 Dec 22;276(1677):4261-70. doi: 10.1098/rspb.2009.0896. |
| Ryan JF, Pang K, Schnitzler CE, Nguyen AD, Moreland RT, Simmons DK, Koch BJ, Francis WR, Havlak P; NISC Comparative Sequencing Program, Smith SA, Putnam NH, Haddock SH, Dunn CW, Wolfsberg TG, Mullikin JC, Martindale MQ, Baxevanis AD. The genome of the ctenophore Mnemiopsis leidyi and its implications for cell type evolution. Science. 2013 Dec 13;342(6164):1242592. doi: 10.1126/science.1242592 |
| Moroz LL, Kocot KM, Citarella MR, Dosung S, Norekian TP, Povolotskaya IS, Grigorenko AP, Dailey C, Berezikov E, Buckley KM, Ptitsyn A, Reshetov D, Mukherjee K, Moroz TP, Bobkova Y, Yu F, Kapitonov VV, Jurka J, Bobkov YV, Swore JJ, Girardo DO, Fodor A, Gusev F, Sanford R, Bruders R, Kittler E, Mills CE, Rast JP, Derelle R, Solovyev VV, Kondrashov FA, Swalla BJ, Sweedler JV, Rogaev EI, Halanych KM, Kohn AB. The ctenophore genome and the evolutionary origins of neural systems. Nature. 2014 Jun 5;510(7503):109-14. doi: 10.1038/nature13400 |
| Chang ES, Neuhof M, Rubinstein ND, Diamant A, Philippe H, Huchon D, Cartwright P. Genomic insights into the evolutionary origin of Myxozoa within Cnidaria. Proc Natl Acad Sci U S A. 2015 Dec 1;112(48):14912-7. doi: 10.1073/pnas.1511468112 |
| Whelan NV, Kocot KM, Moroz LL, Halanych KM. Error, signal, and the placement of Ctenophora sister to all other animals. Proc Natl Acad Sci U S A. 2015 May 5;112(18):5773-8. doi: 10.1073/pnas.1503453112 |
| Torruella G, de Mendoza A, Grau-Bové X, Antó M, Chaplin MA, del Campo J, Eme L, Pérez-Cordón G, Whipps CM, Nichols KM, Paley R, Roger AJ, Sitjà-Bobadilla A, Donachie S, Ruiz-Trillo I. Phylogenomics Reveals Convergent Evolution of Lifestyles in Close Relatives of Animals and Fungi. Curr Biol. 2015 Sep 21;25(18):2404-10. doi: 10.1016/j.cub.2015.07.053 |
| Borowiec ML, Lee EK, Chiu JC, Plachetzki DC. Extracting phylogenetic signal and accounting for bias in whole-genome data sets supports the Ctenophora as sister to remaining Metazoa. BMC Genomics. 2015 Nov 23;16:987. doi: 10.1186/s12864-015-2146-4 |
| Arcila D, Ortí G, Vari R, Armbruster JW, Stiassny MLJ, Ko KD, Sabaj MH, Lundberg J, Revell LJ, Betancur-R R. Genome-wide interrogation advances resolution of recalcitrant groups in the tree of life. Nat Ecol Evol. 2017 Jan 13;1(2):20. doi: 10.1038/s41559-016-0020 |
| Whelan NV, Kocot KM, Moroz TP, Mukherjee K, Williams P, Paulay G, Moroz LL, Halanych |

| |
|---|
| KM. Ctenophore relationships and their placement as the sister group to all other animals. Nat Ecol Evol. 2017 Nov;1(11):1737-1746. doi: 10.1038/s41559-017-0331-3 |
| Shen XX, Hittinger CT, Rokas A. Contentious relationships in phylogenomic studies can be driven by a handful of genes. Nat Ecol Evol. 2017 Apr 10;1(5):126. doi: 10.1038/s41559-017-0126 |
| Laumer CE, Fernández R, Lemer S, Combosch D, Kocot KM, Riesgo A, Andrade SCS, Sterrer W, Sørensen MV, Giribet G. Revisiting metazoan phylogeny with genomic sampling of all phyla. Proc Biol Sci. 2019 Jul 10;286(1906):20190831. doi: 10.1098/rspb.2019.0831 |
| Jeon Y, Park SG, Lee N, Weber JA, Kim HS, Hwang SJ, Woo S, Kim HM, Bhak Y, Jeon S, Lee N, Jo Y, Blazyte A, Ryu T, Cho YS, Kim H, Lee JH, Yim HS, Bhak J, Yum S. The Draft Genome of an Octocoral, Dendronephthya gigantea. Genome Biol Evol. 2019 Mar 1;11(3):949-953. doi: 10.1093/gbe/evz043 |
| Kim HM, Weber JA, Lee N, Park SG, Cho YS, Bhak Y, Lee N, Jeon Y, Jeon S, Luria V, Karger A, Kirschner MW, Jo YJ, Woo S, Shin K, Chung O, Ryu JC, Yim HS, Lee JH, Edwards JS, Manica A, Bhak J, Yum S. The genome of the giant Nomura's jellyfish sheds light on the early evolution of active predation. BMC Biol. 2019 Mar 29;17(1):28. doi: 10.1186/s12915-019-0643-7 |
| Erives A, Fritzsch B. A Screen for Gene Paralogies Delineating Evolutionary Branching Order of Early Metazoa. G3 (Bethesda). 2020 Feb 6;10(2):811-826. doi: 10.1534/g3.119.400951 |
| Pandey A, Braun EL. Phylogenetic Analyses of Sites in Different Protein Structural Environments Result in Distinct Placements of the Metazoan Root. Biology (Basel). 2020 Mar 28;9(4):64. doi: 10.3390/biology9040064 |
| Li Y, Shen XX, Evans B, Dunn CW, Rokas A. Rooting the Animal Tree of Life. Mol Biol Evol. 2021 Sep 27;38(10):4322-4333. doi: 10.1093/molbev/msab170 |

**Supplementary Table 2.** Probe sequences used for whole mount in situ hybridization:

| | |
|---|---|
| INXA<br>(GC:<br>39.45%) | ATGTTATTGG AGATATTAGC GAACTTCAAA GGAGCGACAC CTTTCAAAGA AATAGTTCTA<br>GATGACAAGT GGGACCAGAT TAACCGATGT TACATGTTCC TGCTGTGTGT GATTTTCGGA<br>ACTGTCGGAC TGCTACCGGA AGATGCGCCT CCCTGTCTCT CCAGACGATT AGTGTCTGGT<br>GGAAGAATAG AATGTCCTCC TGCTGACCTG TACTTGGAAC CAACAAGGGT TCATCACACA<br>TGGTATCAGT GGATACCGTT TTACTTTTGG GTCATATCCA TAGCGTTCAT TGGTCCTTAC<br>ATAGTCTACA AGCAGCTGGG TGTCAACGAA CTGAAGCCTA TTCTGGCAAT GCTTCATAAC<br>CCGGTTGATG GTGACGATGT TACAAAGGAT CAAATAAGCA AAGTCTCAAG ATGGTTAGCT<br>ATCAAGCTGA ACATCTTTAT CCAAGAAAAA TCTACCTATG CCAAGATCAC TCAGAGCCAT<br>AGGATGTTTA TTCTAATCTT TTTAACTAAA ATATTCTATC TTGGAGTAAG TTTGGCTACA<br>ATGTATTTTA CTGACACCAT GTTTGAATCC GGCCGCTACC TTACTTACGG CAGCGAATGG<br>TTCGCATCTC TCGATAAGCA ATCAAACTAC ACAAGTTTTG TGCGAGACAG ACTGTTCCCG<br>AAAATGGTGG CATGTGAGAT CAAGAGATGG GGTCCTTCAG GTATGGAGGA GGAACAAGGG<br>ATGTGTGTTC TTGCTCCGAA CGTGATGAAC CAATATCTCT TCCTCATCTT CTGGTTCGCA<br>CTCGTCTTCA CCATCTTCTC CAACACCTTC TCCATCTTCT TCTCCGTTTC GACCCACTGT<br>TTTATTGACG GTGGGTACCA GAGGTTTATC CAGAGCTGCT TTCTAAAAGA AAACAGCAAA<br>CTGAAGTTCA TCTATTTCAA TTGTGGGACT ACTGGCCGGA CTTATCTGCA TCTAATTGCC<br>AAAAACGTTA ACCCTCGGAT TTTTGAACAG CTCATCATCA AACTTAGTGC AGATTTAGTT<br>GAGGAGAAAA ATAAGCAACA CTTAAAGGGG TCAAAGGACA TACTAGTTTG A |
| INXB<br>(GC:<br>45.84%) | ATGGTTATTG ACATCCTCTC CGGTTTTAAG GGGATCACGC CCTTCAAAGG CATCACTTTA<br>GACGATGGAT GGGATCAAAT CAACAGGAGT TTTATGTTCG TTCTGTGCGT TTTAATGGGA<br>ACGGTGGTGA CGGTGCGGCA GTACGCGGGA GGAATTATAT CATGTGACGG CTTCACCAAA<br>TACTCGGGCT CGTTTAGCGA GGATTACTGT TGGACACAGG GTCTTTACAC CATCAAGGAA<br>GCCTATGATC TCCTCACCAT GAATGTGCCT TATCCAGGTG TCATCCCTGA GGACATGCCC<br>ACCTGTATCG AGCGGGAACT TATAAATGGA GGCCGAGTGT CCTGCCCCGA CCCTGAAACA<br>GTCAAACCAC CCACAAGAGT CTACCACTTA TGGTACCAAT GGGTTCCATT CTACTTCTGG<br>CTCGCAGCTG CCGCCTTCTT CTTCCCTTAT CTCATCTACA AACATTTCGG CGTTGGCGAC<br>CTCAAACCCC TCATCCAAAT GCTCCACAAC CCTATTGTTG ACGAAGGTGA CCAGAACTGC<br>ATGGCCGAAA AAGCTAGCAT GTGGCTCTTC TACAAACTGA ACGTTTTCAT GAACGAGAAC<br>ACAATCTTCG CTATCCTGAC TGAAAAACAC CGACTTTTCT TCATTGTTAT GCTCGTCAAG<br>GTGCTCTATC TGATCATCAG TATTCTGGCC CTCTACCTTA CGGACGAGAT GTTCCACATC<br>GGCTCCTTTG TCTCGTACGG GAGCGAGTGG GCGACCTCCC TGCCCGAGGG AGACAACGAG<br>ACGACTCTCG TTAAAGACAA ACTCTTCCCC AAGATGGTCG CTTGTGAGAT CAAACGATGG<br>GGACCTACCG GTCTCGAGGA GGAACAGGGC ATGTGTGTGT TGGCTCCCAA CGTTATCAAC<br>CAATATCTCT TCCTCATCCT CTGGTTCGCC ATTATTTTCT GCATTGCTTG CAACTGTTTG<br>TCCGTCCTCT TCGCCCTTAC AAAGCTGGTC TTCGTCTTGG GCTCCTACAA GAGGCTCCTA<br>GCCAGTGCTT TCCTCAAGGA TGAACTCCAC TACAAACACA TGTTCTTCAA CATTGGTACA<br>AGTGGACGAG TTCTCCTACA AATCGTTGCG ACGAATGTGT CACCGCGGGT CTTCGAGTCC<br>ATCATGGCTA ACCTTGCGAC CAAGTTGATA GCTGAACGTT TGAAGGGAAA CGGCAAAGGT<br>AGCGTCTAG |
| INXC<br>(GC:<br>41.58%) | ATGTTTTGTA TTTTATCTGG TACAATCATG ACCTTTAAAC AGAATTTAGG ATCAATAATA<br>CACTGTATAT CGGATGCAAG AGGCGACGAC AGTTCGTTTG CGGATGCTCA TGCGACATTT<br>GTGCAAGACT ATTGTGCTGC TCAAGGGCTG TACACTTTAA AAGAAGTGTA TGACAAGTCT<br>TGGCCAGATG AAATTCCTTA CCCAGGTATT CTCCAAATGA AAACAATCGG TTGTTTCCCG<br>GGGAGACAGT TCAAAAACGG AACCCCCATC CAGTGCCCGG ACGAGAAAGA TCTGAAACCC |

| | |
|---|---|
| | TTCACAACGG TCTATCATGT CTGGTACATG TTCGTACCGT TCTACTTCTG CGCTGTTGGC ATCGCTTTTT ACTTCCCCTA CACGGTTTTC AGACACCTCA GCGGCATCTA CGACATCAAG CCTATGTTGA ACAGCCTTGC CCTCGACATT GGGGCCTACA CGGAGGAGGA CATAAGTCGA CGTATAGACA ATGTCTCGAG GTGGTTGTAC ATCAAGTTGG ATCCCTACAT GAACAACATG CTTCCTTATA CTCAGATAGT TCACAAACAT TCCATCTTTT ACACGGTGAT GTTGGTGAAG GTGATGTACC TAGCTACCAG TGTTTCTATT TTTTACGCCA CTCACCGGAT ATTCGACCAA GGAAACTTTG CACTCTACGG ATACGATGTT CTAATGAGCA TACCACAGGA AACAAGCTAT AAAGTGATGG ACACAATCTT CCCTAAAATG GTTGGCTGTG AGATCAACAT GTGGGGCCGG ACTGGCGAAC AGAGCGAATC TCTTCTGTGT GTCCTCCCTC AAAACATCGG CAACCAATAC TTCTTCCTTA TATTCTGGTT TCTCCTGATT CTCACCATAC TTTCCAACTG TATCTCTGTA ATAGTGACCA TATTCAGATT TATATTCGTT AGTGGGAGCT ACAAAAGGTT CCTGGCTACC AGCCTCTTGA ATCACGAAGA ACGATACAAG CTGGTGTTTA CACATGTCGG CACGACTGGA AGATACATTT TACTGCTCTG TGCCGATCAT AGCAACCCCA AAATATTCGA GGATCTTCTA GAGATCGTCT GTTCCCTTCT CATAGCAAAC TATCACAAAA GAAAGAGGAG TCGGGATAAG GGACACAGTC GAGCGGAGGG GGTAGGGACT AAAGGGCGAC ACGGTCTGTC TTTTGTGGAC TCAACCGTGT GA |
| INXD (GC: 46.80%) | ATGCTGATCT CGAGCTTAGT TCAGTTCAGC AGGTTATCTC CTTTTAAGGA GATAACTATA GATGACGGGT GGGACCAACT TAACAGGAGT TTCATGTTCG TTCTGATGGT TATCTGTGGA ACTATCGTCA CTGTCCGACA ACATACAGGT AACATCATCT CGTGTAACGG TTTCACAAAA TACGACGGAT CCTTCTCCGA GGACTACTGC TGGACGCAGG GACTCTACAC GATCAGGGAG GCGTACCACG TGAGCGACGT CAACGTCCCT TATCCCGGAG TTATCCCGGA GGAGATCCCA CTCTGTCTAG GAGACAATTG TGATAAGCTA GCAAACAGCA ACACCACTCG AGTGTATCAT CTGTGGTACC AGTGGATCCC CTTCTACTTC TGGCTCGCTT CCGCCGCCTT CTTCCTCCCT TATCTGATCT ACAAGAGATA CGGATTTGGA GATATCAAGC CTCTGATCCA CATGCTGTAC AATCCTCTCG ACGGGGACGA AGGAGTGAAG GCAGATTCGG AGAAGGCCTC AATCTGGCTT TATCACAGAT TCTCTATCTA CATGAACGAG CATTCCATGT ACGCCAACTT TATGGAGAGA CACGGAATCG GCATTCTCGT TATCGCTATC AAGGTGATGT ACCTGATCAT CTCCGTCCTA CTCATGGTCA TGACCGCCAT GATGTTCGAG CTGGCTGACT TCAAGCAGTA CGGTATTGTG TGGGCCCAAC AGTGGCCTGA CCCTCCTGCC AATGTCACAG GAATCAAGGA CCTGCTCTTC CCCAAGATGG TTGCTTGCGA GATCAAGAGA TGGGGACCTA CTGGTCTGGA GGACGAGAAC GGAATGTGTG TCCTGGCCCC CAACGTCATC AACCAGTACA TATTCCTCAT CCTCTGGTGG GCCCTTGTTT TCACCATTGT CTCTAACGTT TTCAACGTAC TGGCTGGAGT TATAAGAATC GTCTTCATCT ATGGTTCTTA CCGCCGGATG TTGGCTAGCG CTTTCCTCAG AGATGATCCT CATTACAAGA AGGTCTACTA CAAGATCGGC ACCTCCGGTC GGGTTATCCT GAACATGCTG GCAGCCTCCA TCTCTCCGAC CTGCTTCCAG GAGATCATGA ACAACGTCTG TCCGCGTCTC ATCCGGGCCC ACGTCTCCAA GAAGGGACGA AACCTGGGCG ACGACCCCCT GTTGTAG |

**Supplementary Table 3. Names/accessions of ctenophore innexins**

| FAMILY | *M. leidyi* | *B. ovata* | *P. bachei* | *H. californensis* |
|---|---|---|---|---|
| **INXA** | ML25993a | | | Hcv1.av93.c10.g19.i1 |
| **INXB** | ML25997a | Bova4.221718.t1 | Pbac.3479642 | Hcv1.av93.c10.g246.i1 |
| **INXC** | ML25998a | Bova4.221717.t1 | Pbac.3473233 | Hcv1.av93.c10.g247.i1 |
| **INXD** | ML25999a | Bova4.221716.t1 | Pbac.3463885 | Hcv1.av93.c10.g248.i1 |
| **INXE** | | | Pbac.3471871 | Hcv1.av93.c8.g510.i1 |
| **INXF** | | | | **F1:** Hcv1.av93.c9.g355.i1 |
| | | | | **F2:** Hcv1.av93.c9.g357.i1 |
| **INXG** | **G1:**ML32831a | Bova4.100819.t1 | | Hcv1.av93.c10.g192.i1 |
| | **G2:**ML47742a | | | |
| **INXH** | ML218922a | Bova4.183840.t1 | | Hcv1.av93.c12.g446.i1 |
| **INXJ** | ML129317a | | **J1:**Pbac.3466313 | **J1a:**Hcv1.av93.c6.g103.i1 |
| | | | | **J1b:**Hcv1.av93.c6.g104.i1 |
| | | | **J2:**Pbac.3464407 | **J2:**Hcv1.av93.c1.g815.i1 |
| **INXK** | | Bova4.221914.t1 | | Hcv1.av93.c1.g143.i1 |
| **INXL** | ML07312a | Bova4.303225.t1 | Pbac.3465668 | Hcv1.av93.c3.g73.i1 |
| **INXM** | ML223536a | Bova4.301137.t1 | Pbac.346454 | Hcv1.av93.c1.g681.i4 |
| **INXN** | | Bova4.205910.t1 | Pbac.3465979* | Hcv1.av93.c10.g18.i1 |
| **INXO** | ML036514a | Bova4.177823.t1 | | Hcv1.av93.c1.g857.i1 |
| **INXP** | ML078817a | Bova4.007720.t1 | | |
| **INXQ** | | Bova4.11432.t1 | | Hcv1.av93.c12.g447.i1 |
| **INXR** | | Bova4.00381.t1 | | Hcv1.av93.c13.g362.i1 |

**Supplementary Table 4. Counts of innexin expression in metacells from Sebé-Pedrós (2018) single cell RNA-Seq data**

| **INX ID** | **ML2.2 ID** | **ClusterID (cell type): # cells expressing (% cells in cluster expressing innexin)** |
|---|---|---|
| INXA | ML25993a | C52 (colloblastA): 37 (52%)<br>C26 (unidentified): 4 (02%)<br>C49 (comb cells): 3 (01%)<br>C55 (burkhardt neuron): 3 (04%)<br>C27 (burkhardt neuron): 3 (02%)<br>C31 (burkhardt neuron*): 2 (03%)<br>C24 (epithelial): 2 (02%)<br>C4 (digestive): 2 (02%)<br>C17 (epithelial): 2 (02%)<br>C48 (comb cells): 2 (02%)<br>C41 (unidentified): 1 (01%)<br>C47 (muscle): 1 (01%)<br>C43 (muscle): 1 (00%) |

| | | |
|---|---|---|
| | | C5 (digestive): 1 (01%)<br>C25 (epithelial): 1 (00%)<br>C32 (burkhardt neuron): 1 (01%)<br>C39 (burkhardt neuron*): 1 (01%)<br>C53 (colloblastB): 1 (01%)<br>C3 (digestive): 1 (00%)<br>C42 (unidentified): 1 (01%)<br>C15 (epithelial): 1 (01%)<br>C21 (epithelial): 1 (00%)<br>unassigned (unidentified): 2 |
| INXB | ML25997a | C3 (digestive): 117 (76%)<br>C28 (burkhardt neuron): 102 (64%)<br>C25 (epithelial): 99 (86%)<br>C21 (epithelial): 95 (68%)<br>C27 (burkhardt neuron): 94 (77%)<br>C26 (unidentified): 94 (65%)<br>C7 (digestive): 93 (67%)<br>C43 (muscle): 87 (46%)<br>C14 (epithelial): 84 (72%)<br>C29 (burkhardt neuron): 82 (66%)<br>C4 (digestive): 74 (86%)<br>C17 (epithelial): 69 (88%)<br>C23 (epithelial): 69 (66%)<br>C13 (epithelial): 67 (76%)<br>C18 (epithelial): 60 (86%)<br>C42 (unidentified): 58 (57%)<br>C41 (unidentified): 52 (54%)<br>C44 (unidentified): 50 (76%)<br>C2 (digestive): 48 (39%)<br>C24 (epithelial): 48 (60%)<br>C1 (digestive): 43 (27%)<br>C34 (burkhardt neuron): 43 (41%)<br>C16 (epithelial): 39 (73%)<br>C15 (epithelial): 38 (61%)<br>C6 (digestive): 37 (56%)<br>C9 (digestive): 36 (43%)<br>C20 (epithelial): 35 (76%)<br>C19 (epithelial): 35 (79%)<br>C33 (burkhardt neuron): 35 (25%)<br>C36 (burkhardt neuron): 33 (52%)<br>C8 (digestive): 33 (39%)<br>C47 (muscle): 33 (44%)<br>C40 (burkhardt neuron): 32 (35%)<br>C38 (unidentified): 27 (35%)<br>C49 (comb cells): 27 (17%)<br>C52 (colloblastA): 26 (37%)<br>C35 (burkhardt neuron): 26 (45%)<br>C55 (burkhardt neuron): 25 (35%)<br>C39 (burkhardt neuron*): 24 (47%) |

| | | |
|---|---|---|
| | | C12 (digestive): 23 (40%)<br>C11 (digestive): 22 (30%)<br>C32 (burkhardt neuron): 22 (39%)<br>C51 (photocytes): 21 (34%)<br>C48 (comb cells): 21 (27%)<br>C10 (digestive): 20 (52%)<br>C53 (colloblastB): 19 (19%)<br>C22 (epithelial): 18 (33%)<br>C5 (digestive): 16 (30%)<br>C37 (unidentified): 16 (32%)<br>C45 (muscle): 15 (17%)<br>C54 (tentacle): 15 (35%)<br>C31 (burkhardt neuron*): 12 (20%)<br>C50 (comb cells): 11 (13%)<br>C46 (muscle): 9 (13%)<br>C30 (burkhardt neuron*): 7 (20%)<br>unassigned (unidentified): 284 (?) |
| INXC | ML25998a | C33 (burkhardt neuron): 8 (05%)<br>C34 (burkhardt neuron): 5 (04%)<br>C35 (burkhardt neuron): 4 (07%)<br>C30 (burkhardt neuron*): 3 (08%)<br>C49 (comb cells): 2 (01%)<br>C16 (epithelial): 2 (03%)<br>C28 (burkhardt neuron): 2 (01%)<br>C25 (epithelial): 2 (01%)<br>C27 (burkhardt neuron): 2 (01%)<br>C14 (epithelial): 1 (00%)<br>C7 (digestive): 1 (00%)<br>C13 (epithelial): 1 (01%)<br>C10 (digestive): 1 (02%)<br>C50 (comb cells): 1 (01%)<br>C48 (comb cells): 1 (01%)<br>C40 (burkhardt neuron): 1 (01%)<br>C26 (unidentified): 1 (00%)<br>unassigned (unidentified): 5 |
| INXD | ML25999a | C3 (digestive): 107 (70%)<br>C26 (unidentified): 107 (74%)<br>C25 (epithelial): 105 (92%)<br>C27 (burkhardt neuron): 98 (80%)<br>C28 (burkhardt neuron): 95 (60%)<br>C21 (epithelial): 94 (67%)<br>C7 (digestive): 92 (66%)<br>C14 (epithelial): 91 (78%)<br>C29 (burkhardt neuron): 78 (62%)<br>C43 (muscle): 78 (41%)<br>C13 (epithelial): 72 (81%)<br>C4 (digestive): 69 (80%)<br>C17 (epithelial): 63 (80%) |

| | | |
|---|---|---|
| | | C44 (unidentified): 57 (87%)<br>C23 (epithelial): 54 (52%)<br>C18 (epithelial): 53 (76%)<br>C2 (digestive): 52 (42%)<br>C42 (unidentified): 49 (49%)<br>C24 (epithelial): 44 (55%)<br>C1 (digestive): 43 (27%)<br>C9 (digestive): 37 (45%)<br>C15 (epithelial): 37 (59%)<br>C6 (digestive): 35 (53%)<br>C20 (epithelial): 34 (73%)<br>C41 (unidentified): 34 (35%)<br>C36 (burkhardt neuron): 32 (50%)<br>C8 (digestive): 32 (38%)<br>C47 (muscle): 29 (39%)<br>C16 (epithelial): 28 (52%)<br>C40 (burkhardt neuron): 28 (31%)<br>C49 (comb cells): 27 (17%)<br>C52 (colloblastA): 26 (37%)<br>C11 (digestive): 26 (36%)<br>C48 (comb cells): 25 (32%)<br>C12 (digestive): 24 (42%)<br>C19 (epithelial): 24 (54%)<br>C55 (burkhardt neuron): 22 (30%)<br>C34 (burkhardt neuron): 22 (21%)<br>C50 (comb cells): 22 (26%)<br>C38 (unidentified): 21 (27%)<br>C33 (burkhardt neuron): 21 (15%)<br>C22 (epithelial): 19 (35%)<br>C51 (photocytes): 18 (29%)<br>C5 (digestive): 18 (34%)<br>C45 (muscle): 18 (21%)<br>C32 (burkhardt neuron): 18 (32%)<br>C39 (burkhardt neuron*): 17 (33%)<br>C54 (tentacle): 15 (35%)<br>C35 (burkhardt neuron): 14 (24%)<br>C10 (digestive): 12 (31%)<br>C37 (unidentified): 12 (24%)<br>C30 (burkhardt neuron*): 10 (28%)<br>C53 (colloblastB): 9 (09%)<br>C46 (muscle): 7 (10%)<br>C31 (burkhardt neuron*): 6 (10%)<br>unassigned (unidentified): 241 |
| INXG.1 | ML32831a | C34 (burkhardt neuron): 57 (54%)<br>C49 (comb cells): 28 (17%)<br>C50 (comb cells): 26 (31%)<br>C33 (burkhardt neuron): 24 (17%)<br>C48 (comb cells): 19 (24%)<br>C47 (muscle): 14 (18%) |

| | | |
|---|---|---|
| | | C18 (epithelial): 8 (11%)<br>C27 (burkhardt neuron): 7 (05%)<br>C21 (epithelial): 6 (04%)<br>C2 (digestive): 4 (03%)<br>C32 (burkhardt neuron): 4 (07%)<br>C40 (burkhardt neuron): 4 (04%)<br>C28 (burkhardt neuron): 4 (02%)<br>C29 (burkhardt neuron): 3 (02%)<br>C35 (burkhardt neuron): 3 (05%)<br>C25 (epithelial): 3 (02%)<br>C1 (digestive): 2 (01%)<br>C46 (muscle): 2 (02%)<br>C20 (epithelial): 2 (04%)<br>C17 (epithelial): 2 (02%)<br>C55 (burkhardt neuron): 2 (02%)<br>C39 (burkhardt neuron*): 2 (03%)<br>C30 (burkhardt neuron*): 2 (05%)<br>C51 (photocytes): 1 (01%)<br>C44 (unidentified): 1 (01%)<br>C45 (muscle): 1 (01%)<br>C24 (epithelial): 1 (01%)<br>C4 (digestive): 1 (01%)<br>C13 (epithelial): 1 (01%)<br>C41 (unidentified): 1 (01%)<br>C14 (epithelial): 1 (00%)<br>C43 (muscle): 1 (00%)<br>C7 (digestive): 1 (00%)<br>C3 (digestive): 1 (00%)<br>C23 (epithelial): 1 (00%)<br>unassigned (unidentified): 32 |
| INXG.2 | ML47742a | C34 (burkhardt neuron): 14 (13%)<br>C33 (burkhardt neuron): 8 (05%)<br>C49 (comb cells): 5 (03%)<br>C47 (muscle): 4 (05%)<br>C9 (digestive): 1 (01%)<br>C12 (digestive): 1 (01%)<br>C55 (burkhardt neuron): 1 (01%)<br>C24 (epithelial): 1 (01%)<br>C28 (burkhardt neuron): 1 (00%)<br>C7 (digestive): 1 (00%)<br>C35 (burkhardt neuron): 1 (01%)<br>C25 (epithelial): 1 (00%)<br>C4 (digestive): 1 (01%)<br>C39 (burkhardt neuron*): 1 (01%)<br>C13 (epithelial): 1 (01%)<br>C17 (epithelial): 1 (01%)<br>C50 (comb cells): 1 (01%)<br>unassigned (unidentified): 6 |

| | | |
|---|---|---|
| INXH | ML218922a | C49 (comb cells): 90 (56%)<br>C48 (comb cells): 55 (71%)<br>C50 (comb cells): 33 (39%)<br>C1 (digestive): 5 (03%)<br>C24 (epithelial): 5 (06%)<br>C13 (epithelial): 5 (05%)<br>C27 (burkhardt neuron): 5 (04%)<br>C7 (digestive): 5 (03%)<br>C25 (epithelial): 4 (03%)<br>C32 (burkhardt neuron): 4 (07%)<br>C42 (unidentified): 4 (04%)<br>C21 (epithelial): 4 (02%)<br>C4 (digestive): 3 (03%)<br>C43 (muscle): 3 (01%)<br>C28 (burkhardt neuron): 3 (01%)<br>C26 (unidentified): 3 (02%)<br>C36 (burkhardt neuron): 2 (03%)<br>C31 (burkhardt neuron*): 2 (03%)<br>C2 (digestive): 2 (01%)<br>C40 (burkhardt neuron): 2 (02%)<br>C41 (unidentified): 2 (02%)<br>C55 (burkhardt neuron): 2 (02%)<br>C33 (burkhardt neuron): 2 (01%)<br>C53 (colloblastB): 2 (02%)<br>C29 (burkhardt neuron): 1 (00%)<br>C38 (unidentified): 1 (01%)<br>C18 (epithelial): 1 (01%)<br>C44 (unidentified): 1 (01%)<br>C47 (muscle): 1 (01%)<br>C6 (digestive): 1 (01%)<br>C16 (epithelial): 1 (01%)<br>C5 (digestive): 1 (01%)<br>C45 (muscle): 1 (01%)<br>C35 (burkhardt neuron): 1 (01%)<br>C19 (epithelial): 1 (02%)<br>C14 (epithelial): 1 (00%)<br>C34 (burkhardt neuron): 1 (00%)<br>C39 (burkhardt neuron*): 1 (01%)<br>C3 (digestive): 1 (00%)<br>C23 (epithelial): 1 (00%)<br>unassigned (unidentified): 38 |
| INXJ | ML129317a | C27 (burkhardt neuron): 16 (13%)<br>C26 (unidentified): 8 (05%)<br>C28 (burkhardt neuron): 6 (03%)<br>C53 (colloblastB): 3 (03%)<br>C30 (burkhardt neuron*): 2 (05%)<br>C25 (epithelial): 1 (00%)<br>C36 (burkhardt neuron): 1 (01%)<br>C32 (burkhardt neuron): 1 (01%) |

| | | |
|---|---|---|
| | | C17 (epithelial): 1 (01%)<br>C14 (epithelial): 1 (00%)<br>C42 (unidentified): 1 (01%)<br>C37 (unidentified): 1 (02%)<br>C2 (digestive): 1 (00%)<br>unassigned (unidentified): 8 |
| INXL | ML07312a | C49 (comb cells): 46 (29%)<br>C48 (comb cells): 39 (50%)<br>C50 (comb cells): 22 (26%)<br>C28 (burkhardt neuron): 3 (01%)<br>C43 (muscle): 2 (01%)<br>C52 (colloblastA): 1 (01%)<br>C7 (digestive): 1 (00%)<br>C18 (epithelial): 1 (01%)<br>C4 (digestive): 1 (01%)<br>C32 (burkhardt neuron): 1 (01%)<br>C41 (unidentified): 1 (01%)<br>C42 (unidentified): 1 (01%)<br>C2 (digestive): 1 (00%)<br>unassigned (unidentified): 21 |
| INXM | ML223536a | C49 (comb cells): 36 (22%)<br>C48 (comb cells): 17 (22%)<br>C50 (comb cells): 15 (18%)<br>C43 (muscle): 3 (01%)<br>C35 (burkhardt neuron): 2 (03%)<br>C25 (epithelial): 2 (01%)<br>C51 (photocytes): 1 (01%)<br>C11 (digestive): 1 (01%)<br>C36 (burkhardt neuron): 1 (01%)<br>C31 (burkhardt neuron*): 1 (01%)<br>C6 (digestive): 1 (01%)<br>C46 (muscle): 1 (01%)<br>C27 (burkhardt neuron): 1 (00%)<br>C30 (burkhardt neuron*): 1 (02%)<br>C15 (epithelial): 1 (01%)<br>C37 (unidentified): 1 (02%)<br>unassigned (unidentified): 10 |
| INXO | ML036514a | C34 (burkhardt neuron): 33 (31%)<br>C33 (burkhardt neuron): 32 (23%)<br>C47 (muscle): 17 (22%)<br>C17 (epithelial): 7 (08%)<br>C46 (muscle): 6 (08%)<br>C45 (muscle): 6 (07%)<br>C35 (burkhardt neuron): 6 (10%)<br>C25 (epithelial): 5 (04%)<br>C55 (burkhardt neuron): 5 (07%)<br>C30 (burkhardt neuron*): 5 (14%) |

| | | |
|---|---|---|
| | | C51 (photocytes): 4 (06%)<br>C18 (epithelial): 4 (05%)<br>C29 (burkhardt neuron): 3 (02%)<br>C48 (comb cells): 3 (03%)<br>C42 (unidentified): 3 (03%)<br>C49 (comb cells): 2 (01%)<br>C2 (digestive): 2 (01%)<br>C6 (digestive): 2 (03%)<br>C24 (epithelial): 2 (02%)<br>C4 (digestive): 2 (02%)<br>C13 (epithelial): 2 (02%)<br>C32 (burkhardt neuron): 2 (03%)<br>C20 (epithelial): 2 (04%)<br>C27 (burkhardt neuron): 2 (01%)<br>C40 (burkhardt neuron): 2 (02%)<br>C14 (epithelial): 2 (01%)<br>C38 (unidentified): 1 (01%)<br>C36 (burkhardt neuron): 1 (01%)<br>C44 (unidentified): 1 (01%)<br>C31 (burkhardt neuron*): 1 (01%)<br>C19 (epithelial): 1 (02%)<br>C41 (unidentified): 1 (01%)<br>C43 (muscle): 1 (00%)<br>C7 (digestive): 1 (00%)<br>C53 (colloblastB): 1 (01%)<br>C50 (comb cells): 1 (01%)<br>C21 (epithelial): 1 (00%)<br>C37 (unidentified): 1 (02%)<br>unassigned (unidentified): 14 |
| INXP | ML078817a | C34 (burkhardt neuron): 4 (03%)<br>C33 (burkhardt neuron): 2 (01%)<br>C39 (burkhardt neuron*): 2 (03%)<br>C30 (burkhardt neuron*): 2 (05%)<br>C28 (burkhardt neuron): 1 (00%)<br>C35 (burkhardt neuron): 1 (01%)<br>C25 (epithelial): 1 (00%)<br>C36 (burkhardt neuron): 1 (01%)<br>C13 (epithelial): 1 (01%)<br>C20 (epithelial): 1 (02%)<br>C41 (unidentified): 1 (01%)<br>unassigned (unidentified): 5 |

**Supplementary Table 5. Counts of occurrences of cells with N innexins in Sebé-Pedrós (2018) single cell RNA-Seq data**

| Number of occurrences of innexins (N) | Count of cells with N innexins (total: 6144) | Percentage of cells with N innexins |
|---|---|---|
| 0 | 2301 | 37.5% |
| 1 | 1724 | 28.1% |
| 2 | 1749 | 28.5% |
| 3 | 271 | 11.7% |
| 4 | 67 | 1.1% |
| 5 | 23 | 0.4% |
| 6 | 5 | 0.1% |
| 7 | 3 | < 0.1% |
| 8 | 0 | 0% |
| 9 | 1 | < 0.1% |

**Supplementary Table 6. Counts of innexin co-expression from Sebé-Pedrós (2018) single cell RNA-Seq data**

| | **INXB(2720)** | **INXJ(51)** | **INXP(22)** | **INXG.2(51)** | **INXG.1(272)** | **INXD(2521)** | **INHX(301)** | **INXC(43)** | **INXM(95)** | **INXL(141)** |
|---|---|---|---|---|---|---|---|---|---|---|
| **INXA(74)** | 28(0.378) | 0(0.000) | 0(0.000) | 0(0.000) | 1(0.014) | 28(0.378) | 2(0.027) | 0(0.000) | 1(0.014) | 1(0.014) |
| **INXL(141)** | 46(0.326) | 1(0.020) | 2(0.091) | 6(0.118) | 44(0.312) | 52(0.369) | 100(0.709) | 3(0.070) | 37(0.389) | |
| **INXM(95)** | 37(0.389) | 0(0.000) | 0(0.000) | 3(0.059) | 27(0.284) | 29(0.305) | 60(0.632) | 3(0.070) | | |
| **INXC(43)** | 23(0.535) | 1(0.023) | 2(0.091) | 5(0.116) | 10(0.233) | 23(0.535) | 5(0.116) | | | |
| **INXH(301)** | 110(0.365) | 1(0.020) | 2(0.091) | 9(0.176) | 72(0.265) | 113(0.375) | | | | |
| **INXD(2521)** | 1761(0.699) | 43(0.843) | 10(0.455) | 20(0.392) | 117(0.430) | | | | | |
| **INXG.1(272)** | 123(0.452) | 1(0.020) | 6(0.273) | 32(0.627) | | | | | | |
| **INXG.2(51)** | 31(0.608) | 0(0.000) | 4(0.182) | | | | | | | |
| **INXP(22)** | 14(0.636) | 0(0.000) | | | | | | | | |
| **INXJ(51)** | 40(0.784) | | | | | | | | | |

**Supplemental Video 1. Contractile activity of the *Mnemiopsis Leidyi* muscle cell.** Activity of the cell was recorded simultaneously with electrophysiological recording (Supplemental Figure 3). Glutamate repetitively evokes the cell contractions. Video - 50 fps. Actual sampling frequency – 4Hz.

**Supplemental File 1. Innexin_GitHub_SnapShot.tar.gz.** Several scripts, alignments, trees, and other files including a phylotocol (DeBiasse and Ryan, 2019) outlining pre-planned phylogenetic analyses are available at: https://github.com/josephryan/ctenophore_innexins This file is a snapshot of that repository made on October 6th before submission.
